# Altered m^6^A modification of specific cellular transcripts affects *Flaviviridae* infection

**DOI:** 10.1101/670984

**Authors:** Nandan S. Gokhale, Alexa B.R. McIntyre, Melissa D. Mattocks, Christopher L. Holley, Helen M. Lazear, Christopher E. Mason, Stacy M. Horner

## Abstract

The RNA modification *N6*-methyladenosine (m^6^A) can modulate mRNA fate and thus affect many biological processes. We analyzed m^6^A modification across the transcriptome following infection by dengue virus (DENV), Zika virus (ZIKV), West Nile virus (WNV), and hepatitis C virus (HCV). We found that infection by these viruses in the *Flaviviridae* family alters m^6^A modification of specific cellular transcripts, including *RIOK3* and *CIRBP*. During viral infection, the addition of m^6^A to *RIOK3* promotes its translation, while loss of m^6^A in *CIRBP* promotes alternative splicing. Importantly, we found that activation of innate immune sensing or the endoplasmic reticulum (ER) stress response by viral infection contributes to the changes in m^6^A modification in *RIOK3* and *CIRBP*, respectively. Further, several transcripts with infection-altered m^6^A profiles, including *RIOK3* and *CIRBP*, encode proteins that influence DENV, ZIKV, and HCV infection. Overall, this work reveals that cellular signaling pathways activated during viral infection lead to alterations in m^6^A modification of host mRNAs to regulate infection.

## Introduction

Transcriptional and post-transcriptional regulation influence gene expression in cells following infection by viruses, including those in the *Flaviviridae* family. The *Flaviviridae* family of positive sense RNA viruses includes dengue virus (DENV), Zika virus (ZIKV), West Nile virus (WNV), and hepatitis C virus (HCV), all of which cause significant mortality and morbidity worldwide. The effects of *Flaviviridae* infection on human health are diverse, ranging from microcephaly and encephalitis to chronic liver disease (Holbrook, 2017; Thrift et al., 2017). Previous studies have shown broad changes in cellular transcript levels during *Flaviviridae* infection that highlight a complex relationship between viral infection and gene expression, whereby the host attempts to resist infection by up- or down-regulating relevant genes while viruses co-opt host transcription to facilitate replication and avoid host defenses (Fink et al., 2007; Kumar et al., 2016; Rosenberg et al., 2018; Sessions et al., 2013; Su et al., 2002; Zanini et al., 2018). Differential expression of proviral and antiviral host factors is therefore an important determinant of the outcome of *Flaviviridae* infection

Host gene expression during *Flaviviridae* infection can be tuned by post-transcriptional RNA controls (De Maio et al., 2016; Luna et al., 2015; Schwerk et al., 2015). An important post-transcriptional RNA regulatory mechanism is the chemical modification of RNA (Gilbert et al., 2016). The most prevalent internal modification of mRNA is *N6*-methyladenosine (m^6^A). The m^6^A epitranscriptome is controlled by specific cellular proteins. METTL3, METTL14, WTAP, and other “writer” proteins form a complex that catalyzes the methylation of adenosine residues in mRNA.

This protein complex targets the consensus motif DRA*CH (where D=G/A/U, R=G/A, H=U/A/C, and * denotes modified A) in mRNA for methylation, although how specific DRACH motifs are selected for modification is still not well understood (Meyer and Jaffrey, 2017; Shi et al., 2019; Yang et al., 2018). “Reader” RNA-binding proteins recognize m^6^A to modulate many aspects of mRNA metabolism, including mRNA splicing, nuclear export, stability, translation, and structure (Meyer and Jaffrey, 2017; Shi et al., 2019; Yang et al., 2018). By regulating specific transcripts, m^6^A plays a role in many important biological processes including circadian rhythm, cell differentiation, development, stress responses, cancer, and viral infection (Gonzales-van Horn and Sarnow, 2017; Meyer and Jaffrey, 2017; Shi et al., 2019; Yang et al., 2018).

Viral infection can be affected by m^6^A modification of either viral or host transcripts. Transcripts from both DNA and RNA viruses can be methylated, and m^6^A in these RNAs has been shown to have various proviral and antiviral functions (Courtney et al., 2017; Gokhale and Horner, 2017; Gokhale et al., 2016; Hao et al., 2019; Imam et al., 2018; Kennedy et al., 2016; Lichinchi et al., 2016a; Lichinchi et al., 2016b; McIntyre et al., 2018; Rubio et al., 2018; Tirumuru et al., 2016; Tsai et al., 2018; Winkler et al., 2019; Ye et al., 2017). m^6^A in specific cellular transcripts is also important during viral infection. For example, m^6^A regulates the expression of the antiviral *IFNB1* transcript induced by double-stranded DNA viruses (Rubio et al., 2018; Winkler et al., 2019). However, the role of m^6^A in cellular mRNA during viral infection is still not well understood, in part because of difficulties in accurately and quantitatively mapping the modification. While ZIKV, Kaposi’s sarcoma-associated herpes virus (KSHV), and human immunodeficiency 1 (HIV-1) have been reported to alter m^6^A modification in cellular mRNAs (Hesser et al., 2018; Lichinchi et al.; Lichinchi et al., 2016b; Tan et al., 2018), the scale of these changes has likely been overestimated (McIntyre et al., 2019). Moreover, there are almost no data on common m^6^A changes in host mRNA across multiple viruses, and the functional consequences of epitranscriptomic changes in cellular mRNA during viral infection have also not been examined. Therefore, better estimating the number of m^6^A changes and defining the consequences of altered m^6^A modification of cellular mRNA during viral infection are important for understanding post-transcriptional regulation of the host response to infection.

Here, we studied the effect of DENV, ZIKV, WNV, and HCV infection on the m^6^A epitranscriptome. We found that infection by all four viruses led to altered m^6^A modification of a set of specific cellular transcripts, and that activation of cellular pathways, including innate immunity and endoplasmic reticulum (ER) stress responses, by infection contribute to differential m^6^A modification and changes in translation or splicing of these transcripts. Importantly, transcripts with altered m^6^A encode proteins that regulate infection, indicating that post-transcriptional gene regulation of mRNA by m^6^A has the potential to affect viral replication.

## Results

### *Flaviviridae* infection alters m^6^A modification of specific cellular transcripts

*Flaviviridae* infection leads to changes in the expression of proviral and antiviral gene products (Fink et al., 2007; Kumar et al., 2016; Rosenberg et al., 2018; Sessions et al., 2013; Su et al., 2002; Zanini et al., 2018). Since m^6^A can modulate RNA fate, and therefore protein expression, we hypothesized that altered m^6^A modification would influence expression of host genes that regulate viral infection. We therefore sought to measure changes in the m^6^A modification of host transcripts during *Flaviviridae* infection using methylated RNA immunoprecipitation and sequencing (MeRIP-seq) (Figure 1A). For MeRIP-seq, we used an anti-m^6^A antibody to enrich m^6^A-modified RNA fragments prior to RNA sequencing of both the input and immunoprecipitated (IP) fractions (Dominissini et al., 2012; Meyer et al., 2012). We note that this antibody can also recognize the similar modification *N6*,2’-O-dimethyladenosine (m^6^A_m_), which is found at lower abundance than m^6^A and, in mRNA, only within the 5’ cap (Linder et al., 2015; Mauer and Jaffrey, 2018). We performed MeRIP-seq on RNA extracted from human Huh7 liver hepatoma cells, which are permissive for infection by all four viruses. At 48 hours post-infection, 60-90% of cells were infected with DENV, ZIKV, WNV, or HCV, depending on the virus (Figure S1A). We first identified the gene expression changes in response to infection. To do this, we analyzed differential expression of genes between infected samples and uninfected controls using the input fractions from MeRIP-seq and found 50 genes that were significantly differentially expressed (DESeq2, adjusted p < 0.05, |Log_2_Fold Change (FC)| ≥ 2) following infection by all four viruses individually (Figure S1B-C, Table S1). Notably, although ZIKV, DENV, and WNV are known to generate acute, cytotoxic responses, while HCV leads to persistent infection (Neufeldt et al., 2018), we found that several pathways were similarly altered by all of these viruses in Huh7 cells. Significantly upregulated pathways included those associated with innate immunity (such as NF-κB, TNF, and MAPK signaling) and the ER stress response, while downregulated pathways included those associated with the cell cycle (Figure S1D). These results, which we validated by RT-qPCR, are similar to what has been reported for individual *Flaviviridae* (Figure S1E) (Fink et al., 2007; Kumar et al., 2016; Rosenberg et al., 2018; Sessions et al., 2013; Su et al., 2002; Zanini et al., 2018).

**Figure 1:**
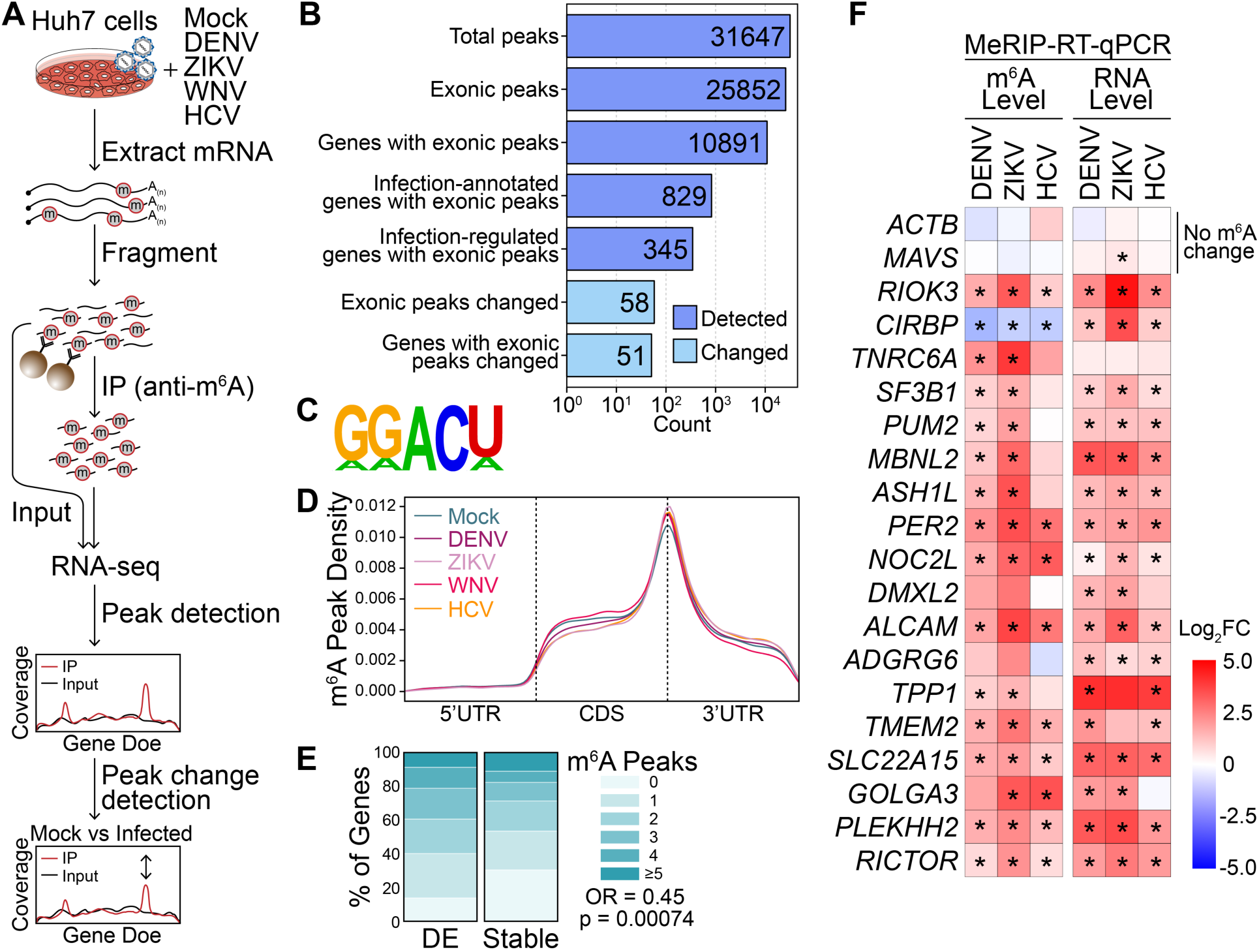
*Flaviviridae* infection alters m^6^A modification of specific transcripts. **(A)** Schematic of the MeRIP-seq protocol used to identify differential m^6^A methylation following infection of Huh7 cells with DENV, ZIKV, WNV, and HCV. RNA was harvested at 48 hours post-infection (hpi) and experiments were performed in triplicate. **(B)** The number of peaks and genes with m^6^A peaks detected in ≥ 2 mock- or virus-infected samples (dark blue; MACS2 q-value < 0.05) and peaks that change during infection (light blue, |peak – gene Log_2_FC| ≥ 1, adjusted p < 0.05). “Infection-annotated genes” are defined as those with known annotations for the Reactome Pathways ‘Infectious Disease’, ‘Unfolded Protein Response’, ‘Interferon Signaling’, or ‘Innate Immune Signaling’ in the database used by fgsea. “Infection-regulated genes” are defined as those that show a Log_2_ fold change in gene expression ≥ 2 in RNA expression between mock- and virus-infected samples in our data set (adjusted p < 0.05). **(C)** The most significantly enriched motif identified as enriched in the MeRIP fractions across all samples (HOMER, p = 1e-831). **(D)** Metagene plot of methylated DRACH motifs across transcripts in mock- and virus-infected cells. DRACH motifs were considered methylated if they fell under a peak detected in at least two replicates. **(E)** The percent of genes with m^6^A peaks that changed expression with infection (|Log_2_FC| ≥ 2, adjusted p < 0.05, N = 137) and genes that remained stable (|Log_2_FC| < 0.5, adjusted p > 0.05, N = 7627) for transcripts with mean expression ≥ 50 reads. **(F)** (Left) MeRIP-RT-qPCR analysis of relative m^6^A level of transcripts with infection-altered m^6^A modification or controls (*ACTB and MAVS*) in DENV, ZIKV, and HCV-infected (48 hpi) Huh7 cells. (Right) RNA expression of these transcripts relative to *GAPDH* (right). Values in heatmaps are the mean of 3 independent experiments. * p < 0.05, by unpaired Student’s t test.

We then predicted m^6^A-modified regions by calling peaks in IP over input RNA-seq coverage across transcripts using MACS2 (Zhang et al., 2008). We detected a total of 31,647 peaks, with 25,852 exonic peaks corresponding to 10,891 genes across all uninfected and infected samples (Figure 1B). The known m^6^A motif DRACH (in particular, GGACU), was enriched under the identified peaks (Figure 1C). As expected, detected peaks were most common at the end of the coding sequence and beginning of the 3’ untranslated region (UTR) (Figure 1D) (Meyer and Jaffrey, 2017; Shi et al., 2019; Yang et al., 2018). We did not observe a change in the distribution of m^6^A across transcript regions with DENV, ZIKV, WNV, or HCV infection (Figure 1D). This is in contrast to a previous report that suggested ZIKV infection was associated with increased methylation at the 5’ UTRs of cellular transcripts (Lichinchi et al., 2016b); however, we also did not detect a difference in m^6^A distribution following ZIKV infection on reanalysis of that published data (Figure S1F). Further, following viral infection, we found only subtle changes in the overall level of m^6^A relative to unmodified adenosine in purified mRNA, as analyzed by liquid chromatography tandem-mass spectrometry (LC-MS/MS) of digested nucleotides, and no change in the expression of cellular m^6^A machinery, as analyzed by immunoblotting (Figure S1G-H). Indeed, since the expression of the methylation machinery was not changed by infection, we would not predict broad, unidirectional changes in the abundance or distribution of m^6^A on cellular transcripts.

However, functional annotation of the m^6^A-modified genes expressed in the infected samples did reveal an enrichment for genes with roles in infection. In total, 829 methylated genes were annotated as involved in the Reactome Pathways of “Infectious Disease”, “Unfolded Protein Response”, “Interferon Signaling”, or “Innate Immune System” (“Infection-annotated genes”; see Methods; Figure 1B). Further, 345 genes that were differentially expressed between infected and uninfected samples were also methylated (“Infection-regulated genes”; Figure 1B). Indeed, mRNAs that changed expression with infection (p adj < 0.05, |Log_2_FC| ≥ 2, mean expression ≥ 50) were more likely to have at least one m^6^A site than those that did not change expression (p adj > 0.05, |Log_2_FC| < 0.5, mean expression ≥ 50; Fisher’s exact test p = 0.00074, odds ratio = 0.64) (Figure 1E). These results are consistent with previous reports that genes that undergo dynamic regulation tend to contain more m^6^A sites in their transcripts than stable housekeeping genes (Schwartz et al., 2014), and suggest that m^6^A may be an important regulator of genes implicated in infection.

We next predicted changes in m^6^A based on differences in IP enrichment relative to gene expression with infection by all four *Flaviviridae* members. We detected shared m^6^A changes in 58 exonic peaks in 51 genes following infection with all viruses, most of which showed increases in m^6^A and occurred within the 3’ UTR or coding sequence of the transcript (Figure 1B, Table S2). Whereas genes that showed changes in expression were enriched for pathways with known roles in infection (Figure S1D), genes that showed changes in methylation did not show any enrichment for functional categories relevant to infection. We and others previously showed that MeRIP-RT-qPCR can detect relative changes in m^6^A levels (Engel et al., 2018; McIntyre et al., 2019). Therefore, we used this method with RT-qPCR primers under the changed m^6^A peaks to orthogonally validate a set of 18 of the predicted m^6^A changes in transcripts following infection. In these and subsequent analyses, we focused on m^6^A changes following infection by DENV, ZIKV, and HCV. Of the 18 transcripts tested by MeRIP-RT-qPCR, 16 showed a significant change in m^6^A modification relative to any change in gene expression with at least two viruses, and 9 of those showed a significant change with all three viruses. The control mRNAs (*ACTB* and *MAVS*), both predicted to be stably methylated during infection, indeed showed no m^6^A changes (Figure 1F). Most non-significant m^6^A changes trended towards the change predicted by MeRIP-seq (Figure 1F).

For our predictions of pan-viral m^6^A changes using MeRIP-seq (above), we compared all infected to all uninfected replicates for increased statistical power (McIntyre et al., 2019). However we also wanted to detect any peak changes unique to single viruses, and therefore, we used the same computational approach described above to identify significant peaks unique to each virus (Table S2). MeRIP-RT-qPCR validation of these putative virus-specific peaks (two per virus) showed similar changes in relative m^6^A modification at those peaks with infection by all three viruses tested, rather than individual virus-mediated changes (Figure S1I), suggesting that most m^6^A regulation occurs through processes activated in response to infection by all the *Flaviviridae* we tested. Together, our data reveal that hundreds of transcripts differentially expressed during *Flaviviridae* infection contain m^6^A and that infection alters m^6^A modification of specific host transcripts.

### *Flaviviridae* infection alters m^6^A modification of *RIOK3* and *CIRBP* mRNA through distinct pathways

We focused on two specific transcripts that gain or lose m^6^A during infection by all viruses (DENV, ZIKV, WNV, and HCV) for further analysis: *RIOK3* (gains) and *CIRBP* (loses). *RIOK3* encodes a serine/threonine kinase and has been implicated in regulating antiviral signaling (Feng et al., 2014; Takashima et al., 2015; Willemsen et al., 2017), while *CIRBP* encodes a stress-induced RNA-binding protein (Liao et al., 2017). Following viral infection, *RIOK3* mRNA gains an m^6^A peak in the 3’ UTR close to the stop codon (Figure 2A), and *CIRBP* mRNA shows reduced m^6^A modification in the coding sequence of its last exon (Figure 2B). The *RIOK3* and *CIRBP* peaks span four and three DRACH motifs, respectively. Both peaks have been previously reported in published datasets, although the function of m^6^A on these transcripts has not been investigated; the *RIOK3* peak was identified in mouse liver tissue (Zhou et al., 2018), while the CIRBP peak was present in HepG2 cells (Huang et al., 2019; Zhong et al., 2018). We performed MeRIP-RT-qPCR on RNA from cells infected with DENV, ZIKV, and HCV to validate these predicted changes in m^6^A, as in Figure 1F. MeRIP-RT-qPCR confirmed that relative to gene expression, the m^6^A modification of *RIOK3* significantly increased following infection and that of *CIRBP* decreased, while *RIOK3* and *CIRBP* mRNA levels both increased following infection (Figure 1F and 2C). We found similar changes in the m^6^A modification of *RIOK3* and *CIRBP* in chromatin-associated RNA following ZIKV infection, suggesting that the regulation of m^6^A at these sites occurs co-transcriptionally (Ke et al., 2017; Slobodin et al., 2017) (Figure S2A). In uninfected cells, both *RIOK3* and *CIRBP* transcripts are bound by the m^6^A-binding protein YTHDF1 (Figure S2B-C). However, DENV, ZIKV, and HCV infection all increased the association of YTHDF1 with *RIOK3* mRNA and decreased its association with *CIRBP* mRNA, suggesting that YTHDF1 recognizes the altered m^6^A modification of *RIOK3* and *CIRBP* transcripts following infection (Figure S2D).

**Figure 2:**
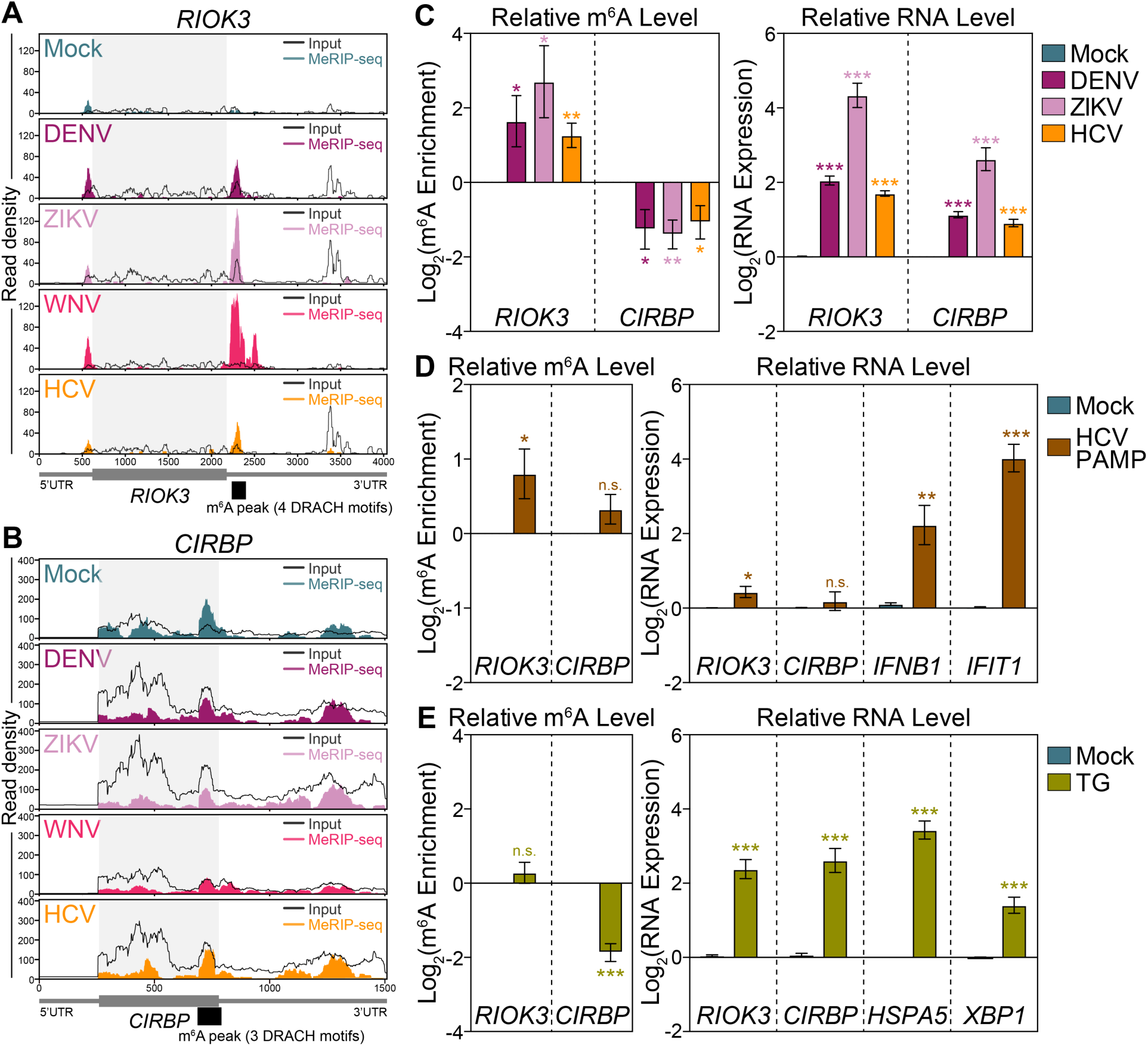
*Flaviviridae* infection alters m^6^A modification of *RIOK3* and *CIRBP* mRNA through distinct cellular pathways. **(A and B)** Coverage plot of MeRIP (color) and input (black) reads in (A) *RIOK3* and (B) *CIRBP* transcripts in Huh7 cells infected with the indicated virus (48 hpi) as determined by MeRIP-seq. Data are representative of three biological replicates. Infection-altered m^6^A peaks (and the number of DRACH motifs within) are indicated in black under the transcript map. **(C)** (Left) MeRIP-RT-qPCR analysis of relative m^6^A level of *RIOK3* and *CIRBP* in mock- and virus-infected (48 hpi) Huh7 cells. (Right) RNA expression of *RIOK3* and *CIRBP* relative to *HPRT1* (right). **(D)** (Left) MeRIP-RT-qPCR analysis of relative m^6^A level of *RIOK3* and *CIRBP* in mock- and HCV PAMP-transfected (8 h) Huh7 cells. (Right) RNA expression of *RIOK3, CIRBP*, as well as positive control transcripts *IFNB1* and *IFIT1* relative to *HPRT1*. **(E)** (Left) MeRIP-RT-qPCR analysis of relative m^6^A level of *RIOK3* and *CIRBP* in mock- and thapsigargin-treated (TG; 16 h) Huh7 cells. (Right) RNA expression of *RIOK3, CIRBP*, and positive control transcripts *HSPA5* and *XBP1* relative to *HPRT1*. Values are the mean ± SEM of 6 (C-D), 3 (E), or 5 (F) biological replicates. * p < 0.05, ** p < 0.01, *** p < 0.001 by unpaired Student’s t test. n.s. = not significant.

We next investigated whether cellular pathways stimulated by viral infection (Figure S1D) contribute to the virally induced m^6^A changes in *RIOK3* and *CIRBP. Flaviviridae* infection drives innate immune signaling cascades which lead to transcriptional induction of interferon-β (IFN) and antiviral interferon-stimulated genes (ISGs) (Horner and Gale, 2013; Munoz-Jordan and Fredericksen, 2010; Suthar et al., 2013). Therefore, we tested whether innate immune activation in the absence of a replicating virus alters m^6^A modification. We measured the relative m^6^A levels of *RIOK3* and *CIRBP* mRNA by MeRIP-RT-qPCR following transfection of Huh7 cells with the short, previously described, HCV pathogen-associated molecular pattern (HCV PAMP) immunostimulatory RNA (Saito et al., 2008). As expected, HCV PAMP induced expression of *IFNB1* and the ISG *IFIT1* (Figure 2D). It also increased m^6^A modification of *RIOK3* mRNA to a similar degree as viral infection, but it did not reproduce the decrease in *CIRBP* methylation seen with viral infection (Figure 2D). These data indicate that innate immune signaling promotes the m^6^A modification of *RIOK3* mRNA following infection.

Our work and that of others have shown that the ER stress response is activated during *Flaviviridae* infection, which remodels host ER membranes to facilitate viral replication (Figure S1D) (Blazquez et al., 2014; Chan, 2014; Neufeldt et al., 2018). The ER Ca^2+^ ATPase inhibitor thapsigargin can induce a similar stress response, including increased expression of *HSPA5* and *XBP1* (Lee et al., 2012). To test whether ER stress alters the m^6^A modification of *RIOK3* and *CIRBP*, we measured their relative m^6^A levels by MeRIP-RT-qPCR following treatment of cells with thapsigargin (Figure 2E). Thapsigargin treatment increased the mRNA level of both *RIOK3* and *CIRBP*, as well as that of the positive controls *HSPA5* and *XBP1*, by about 4-fold. However, while thapsigargin treatment did not change the relative m^6^A level of *RIOK3*, it did reduce m^6^A modification of *CIRBP*, similar to what we observed with viral infection (Figure 2E). Taken together, these data reveal that innate immune and ER stress signaling, which are activated during *Flaviviridae* infection, can separately affect m^6^A modification of different transcripts.

### m^6^A modification enhances RIOK3 protein expression during infection

We next investigated the function of m^6^A in *RIOK3* mRNA during infection. Consistent with our finding that DENV, ZIKV, and HCV infection all increased *RIOK3* mRNA levels (Figure 2C), RIOK3 protein expression also increased following infection (Figure 3A). m^6^A can alter mRNA nuclear export, stability, and translation, all of which could regulate protein expression (Meyer and Jaffrey, 2017; Yang et al., 2018). When we analyzed the mRNA levels of *RIOK3* in nuclear and cytoplasmic fractions from uninfected and infected Huh7 cells using RT-qPCR, we found no significant change in the nuclear export of *RIOK3* post-infection (Figure S3A). Similarly, we found no consistent change in the mRNA stability of *RIOK3* in uninfected and infected cells (Figure S3B). However, we did detect increased nascent translation of RIOK3 in DENV-infected cells compared to uninfected cells as measured by ^35^S labeling of nascent proteins followed by RIOK3 protein immunoprecipitation, suggesting that RIOK3 translation was increased in infected cells (Figure 3B). This is consistent with our observation that during infection *RIOK3* has increased binding to the m^6^A reader protein YTHDF1, which is known to promote translation of bound mRNAs under specific conditions (Figure S2D) (Han et al., 2019; Shi et al., 2018; Wang et al., 2019; Wang et al., 2015). However, global cellular translation is known to be inhibited during DENV, ZIKV, and HCV infection (Arnaud et al., 2010; Garaigorta and Chisari, 2009; Roth et al., 2017). Indeed, we observed that in Huh7 cells, infection with all three viruses induces the phosphorylation of the eukaryotic translation initiation factor eIF2α, which inhibits recycling of eIF2 and therefore prevents translation of RNAs that require this factor (Figure S3C) (Stern-Ginossar et al., 2019; Wek, 2018). Together, our results suggest that m^6^A modification of *RIOK3* could allow this transcript to be efficiently translated during infection, despite global inhibition of translation.

**Figure 3:**
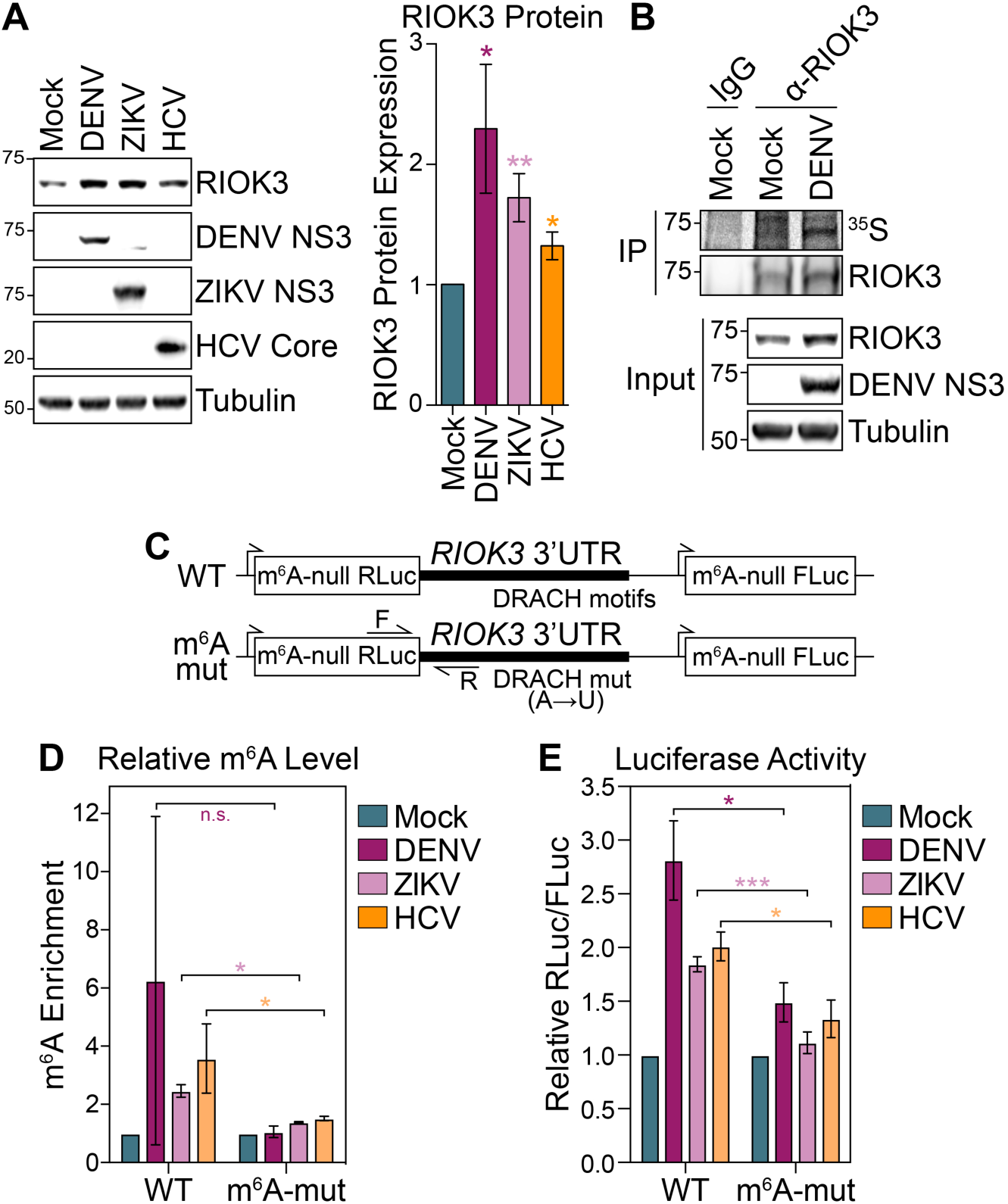
m^6^A promotes RIOK3 protein expression. **(A)** (Left) Representative immunoblot of RIOK3 protein expression in mock- and virus-infected (48 hpi) Huh7 cells. (Right) Quantification of RIOK3 protein expression relative to tubulin from replicate experiments. **(B)** Immunoprecipitation (IP) of RIOK3 from mock- and DENV-infected (48 hpi) Huh7 cells labeled with ^35^S for 3 hours. IP fractions were analyzed by autoradiography (^35^S) and immunoblotting. Data are representative of 3 biological replicates. **(C)** Schematic of WT and mutant m^6^A-null *Renilla* luciferase (RLuc) *RIOK3* 3’ UTR reporters that also express m^6^A-null Firefly luciferase (FLuc) from a separate promoter. RT-qPCR primer (F and R) locations are indicated with arrows. **(D)** MeRIP-RT-qPCR analysis of relative m^6^A level of stably expressed WT and m^6^A-mut *RIOK3* 3’ UTR reporter RNA in mock- and virus-infected (48 hpi) Huh7 cells. **(E)** Relative luciferase activity (RLuc/FLuc) in mock- and virus-infected (48 hpi) Huh7 cells stably expressing WT and m^6^A-mut *RIOK3* 3’ UTR reporters. Relative luciferase activity in uninfected cells was set as 1 for each reporter. Values are the mean ± SEM of 6 (A), 2 (D), or 5 (E) biological replicates. * p < 0.05, ** p < 0.01, *** p < 0.001 by unpaired Student’s t test. n.s. = not significant.

To directly test whether m^6^A can promote RIOK3 protein expression during infection, we generated Huh7 cell lines stably expressing a luciferase reporter which contains the wild type (WT) *RIOK3* 3’ UTR, or an analogous 3’ UTR sequence in which all putative m^6^A sites were abrogated by A*→*T mutations (m^6^A-mut), downstream of a *Renilla* luciferase gene in which all DRACH motifs were ablated (m^6^A-null) (Figure 3C). These constructs also expressed a separate m^6^A-null Firefly luciferase gene whose expression is not regulated by m^6^A. As expected, the WT *RIOK3* reporter had increased m^6^A modification compared to the m^6^A-mut *RIOK3* reporter following viral infection, as measured by MeRIP-RT-qPCR using primers that specifically amplified reporter RNA (Figure 3D). This reveals that the *RIOK3* 3’ UTR sequence is sufficient for m^6^A addition following infection. Importantly, the relative luciferase activity of the WT *RIOK3* reporter was significantly increased compared to the m^6^A-mut reporter following viral infection (Figure 3E). Taken together, these data reveal that m^6^A modification of the 3’ UTR of *RIOK3* mRNA during infection consistently promotes its translation during infection with all three *Flaviviridae* tested.

### m^6^A modification promotes alternative splicing of *CIRBP* mRNA during infection

We then analyzed the function of reduced m^6^A modification in *CIRBP* mRNA following infection. Neither the nuclear export nor the stability of *CIRBP* mRNA were consistently affected following DENV, ZIKV, or HCV infection, suggesting that the loss m^6^A in *CIRBP* does not regulate these processes (Figure S4A-B). Based on our RNA-seq data, *CIRBP* encodes at least 2 isoforms: (1) the dominant, short isoform (*CIRBP-S*) which encodes a 172 aa, 18 kDa protein and (2) a second, long isoform in which an intron immediately downstream of the infection-altered m^6^A peak and upstream of the stop codon is retained (*CIRBP-L*), resulting in a 297 aa, 32 kDa protein (Figure 4A; retained intron referred to as alternatively spliced region (ASR)). Interestingly, analysis of our RNA-seq data using MAJIQ (Vaquero-Garcia et al., 2016) to identify local splice variants suggested decreased retention of this intron during infection, which we confirmed in DENV, ZIKV, and HCV-infected cells using RT-qPCR (Figure 4B). We observed a similar reduction of intron retention following thapsigargin treatment, which we had found also reduces m^6^A modification of *CIRBP* (Figure 4C and 2F). Indeed, both viral infection and thapsigargin treatment significantly reduced the protein level of CIRBP-L containing the retained intron, while not affecting expression of CIRBP-S (Figure 4D-E). To test whether reduction of m^6^A modification at the m^6^A peak in *CIRBP* might affect alternative splicing of this transcript, we generated a splicing reporter wherein the m^6^A-null *Renilla* luciferase gene was fused to the WT genomic sequence of *CIRBP* from exon 5 onwards (WT *CIRBP*) and a corresponding reporter in which the putative m^6^A sites in the identified *CIRBP* m^6^A peak were synonymously mutated (m^6^A-mut *CIRBP*) (Figure 4F). Using RT-qPCR, we found that the m^6^A-mut reporter had reduced intron retention compared to the WT reporter, suggesting that the loss of m^6^A in *CIRBP* regulates its alternative splicing and reduces the expression of the long isoform (Figure 4G).

**Figure 4:**
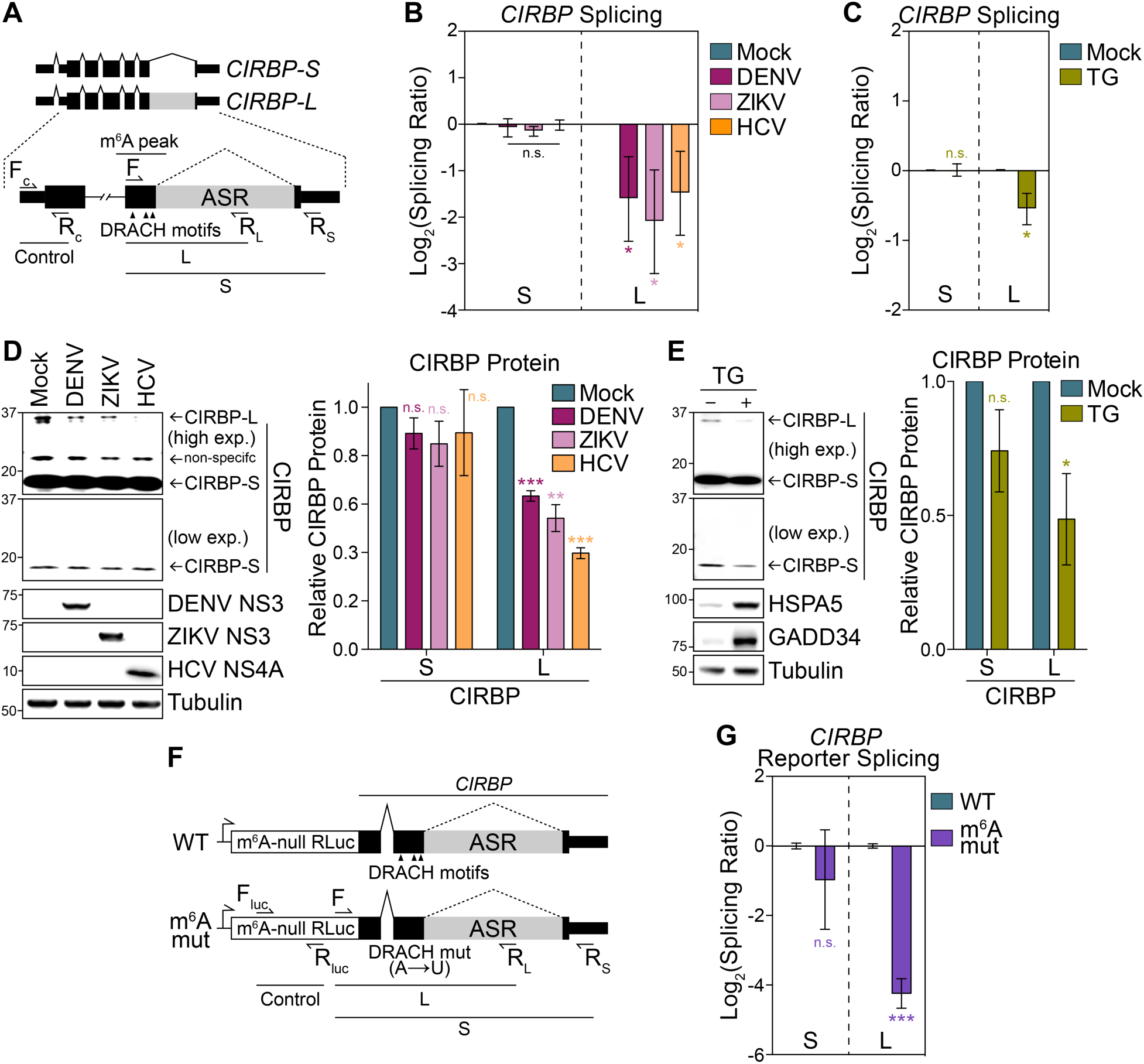
m^6^A promotes alternative splicing of *CIRBP*. **(A)** Schematic of *CIRBP* transcript isoforms with a focus on the alternatively spliced region (ASR). RT-qPCR primer locations are indicated with arrows (F_C_-R_C_: control *CIRBP* amplicon; F-R_L_: long isoform specific; F-R_S_: short isoform specific. **(B)** RT-qPCR analysis of short (S) and long (L) *CIRBP* RNA isoform expression in mock- and virus-infected (48 hpi) Huh7 cells relative to control *CIRBP* amplicon. **(C)** RT-qPCR analysis of S and L *CIRBP* RNA isoform expression in mock- and TG-treated (16 h) Huh7 cells. **(D)** (Left) Representative immunoblot of short (CIRBP-S) and long (CIRBP-L) CIRBP protein isoforms in mock- and virus-infected (48 hpi) Huh7 cells. (Right) Quantification of CIRBP protein isoform expression relative to tubulin from replicate experiments. **(E)** (Left) Representative immunoblot analysis of CIRBP protein isoforms in mock- and TG-treated (500nM, 16 h) Huh7 cells. HSPA5 and GADD34 are positive controls. (Right) Quantification of CIRBP protein isoform expression relative to tubulin from replicate experiments. **(F)** Schematic of WT and m^6^A-mut *CIRBP* splicing reporters. RT-qPCR primer locations (F_luc_-R_luc_: control; F-R_L_: long isoform specific; F-R_s_: short isoform specific) are indicated with arrows. **(G)** RT-qPCR analysis of *CIRBP* splicing reporter isoform expression (S and L) relative to control *RLuc* amplicon in Huh7 cells transfected with WT and m^6^A-mut constructs. Values are the mean ± SEM of 3 (B, D, E, G) or 5 (C) biological replicates. * p < 0.05, ** p < 0.01, *** p < 0.001 by unpaired Student’s t test. n.s. = not significant.

### m^6^A-altered genes regulate *Flaviviridae* infection

Having found that both *RIOK3* and *CIRBP* transcripts have altered m^6^A modification during infection, we next tested whether their encoded protein products affect *Flaviviridae* infection. To this end, we depleted RIOK3 and CIRBP in Huh7 cells using small interfering RNA (siRNA), infected these cells with DENV, ZIKV, or HCV, and then measured viral titer in the supernatant at 72 hours post-infection. siRNA treatment reduced both *RIOK3* and *CIRBP* mRNA levels by ∼70% and did not affect cell viability (Figure S5A). We found that RIOK3 depletion significantly reduced the production of infectious DENV and ZIKV particles but increased the production of infectious HCV particles (Figure 5A). Consistent with these data, RIOK3 stably overexpressed in two different clonal cell lines (RIOK3-1 and RIOK3-2) had the opposite effect on DENV, ZIKV, and HCV infectious particle production (Figure 5B-C). This suggests that RIOK3 promotes DENV and ZIKV infection but inhibits HCV infection. In contrast, the depletion of CIRBP consistently reduced the production of infectious DENV, ZIKV, and HCV (Figure 5D), while overexpression of both the short and long isoforms of CIRBP in two different clonal cell lines (CIRBP-S-1 and 2, CIRBP-L-1 and 2) increased infection by these viruses (Figure 5E-F).

**Figure 5:**
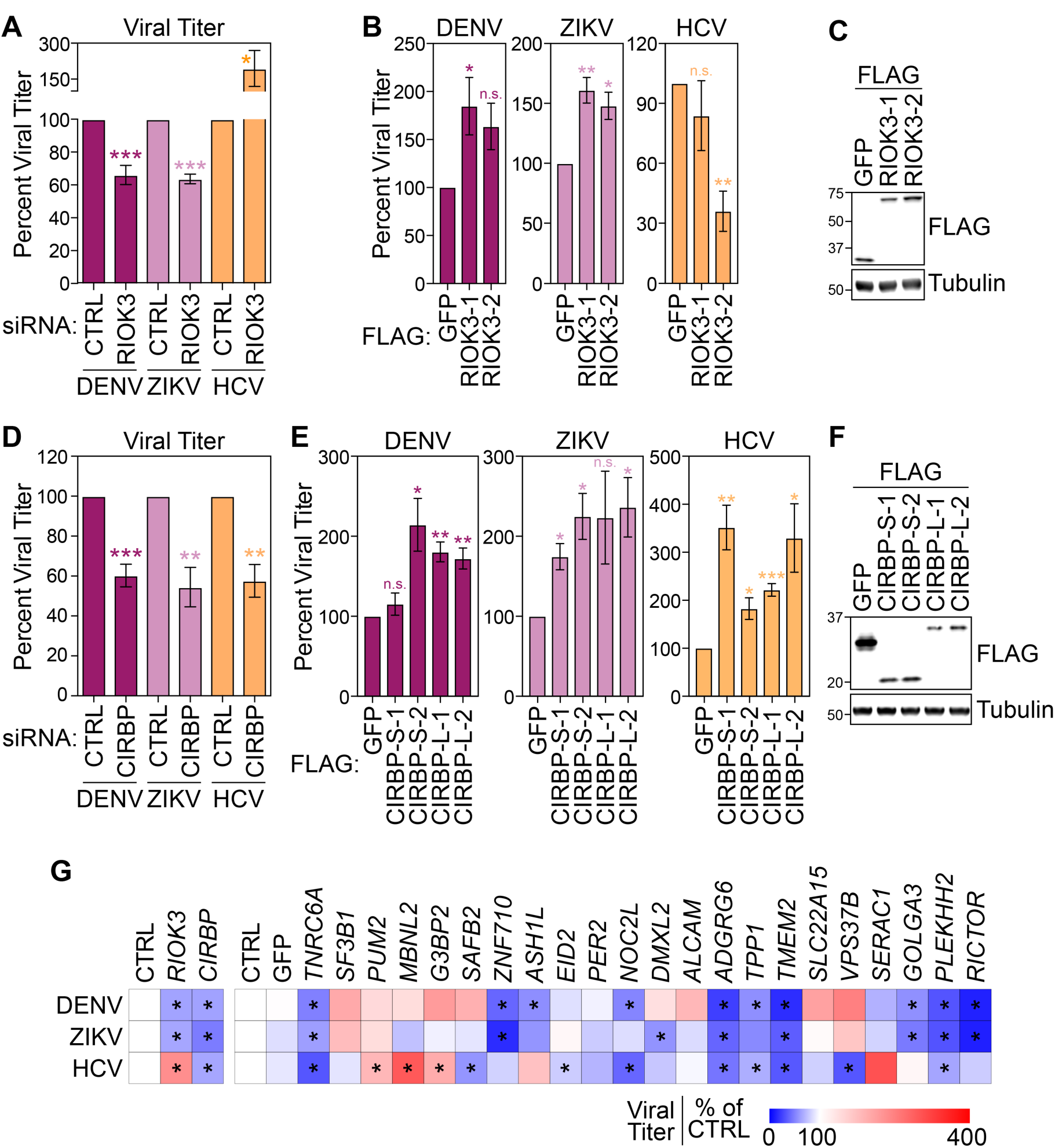
Genes with infection-induced m^6^A alterations regulate *Flaviviridae* infection. **(A)** Focus-forming assay (FFA) of supernatants harvested from DENV, ZIKV, or HCV-infected (72 hpi) Huh7 cells treated with non-targeting control (CTRL) or *RIOK3* siRNA. **(B)** FFA of supernatants harvested from DENV, ZIKV, or HCV-infected (72 hpi) Huh7 cells stably overexpressing FLAG-GFP or FLAG-RIOK3 (2 independent clones). **(C)** Immunoblot analysis of cell lines used in (B). **(D)** FFA of supernatants harvested from DENV, ZIKV, or HCV-infected (72 hpi) Huh7 treated with CTRL or *CIRBP* siRNA. **(E)** FFA of supernatants harvested from DENV, ZIKV, or HCV-infected (72 hpi) Huh7 cells stably overexpressing FLAG-GFP or the short (FLAG-CIRBP-S) or long (FLAG-CIRBP-L) isoforms of CIRBP (2 independent clones). **(F)** Immunoblot analysis of cell lines used in (C). **(G)** Summary of data from targeted siRNA depletion experiments. Viral titers were determined by FFA of supernatants harvested from infected cells (48 hpi) treated with the indicated siRNAs. Data are presented as percentage of titer of each virus relative to cells treated with CTRL siRNA. * indicates significance. Values are the mean ± SEM of 4 (A and D), or 3 (B, E, G) experiments. All viral infections for experiments in this figure were performed at a multiplicity of infection of 0.2. * p < 0.05, ** p < 0.01, *** p < 0.001 by unpaired Student’s t test. n.s. = not significant.

We then performed a targeted siRNA screen to test whether other transcripts with infection-altered m^6^A modification affect *Flaviviridae* infection. We depleted transcripts in which we had identified m^6^A changes during infection (either co-regulated (Figure 1F) or virus-specific (Figure S1I)), infected depleted cells with DENV, ZIKV, or HCV, and measured cell viability, relative RNA depletion levels, and the production of infectious virions in the supernatant at 48 hours post-infection (Figure 5G and S5A-B). We focused only on those transcripts that were depleted by at least 40% in our further analysis (21 out of 24 tested). For these, we found that 86% (18/21) regulate at least 1 virus, while 10/21 affect at least 2, and 6/21 regulate all three viruses. For each virus, ∼50% of m^6^A-altered transcripts that we tested significantly increased or decreased infection. This indicates that by modifying specific transcripts that modulate infection, m^6^A can tune the outcome of infection.

## Discussion

Here, we identify changes in m^6^A methylation of cellular mRNAs during infection by viruses in the *Flaviviridae* family, specifically DENV, ZIKV, WNV, and HCV. We observed that infection by all of these viruses leads to changes in m^6^A modification of a specific set of cellular transcripts, including some that encode factors that modulate *Flaviviridae* infection in Huh7 cells. We found that virus-induced pathways, including innate immune signaling and ER stress responses, contributed to altered m^6^A modification of at least two of these transcripts during infection. Taken together, this work suggests that m^6^A epitranscriptomic changes induced through cellular signaling pathways influence *Flaviviridae* infection.

We identified hundreds of m^6^A-modified transcripts that were differentially expressed during infection or that were annotated as part of cellular pathways relevant for infection. These findings suggest that m^6^A has the potential to post-transcriptionally regulate many genes during infection. Here, we focused on specific transcripts with virus-induced m^6^A changes; we identified 58 peak changes in 51 transcripts following infection by DENV, ZIKV, WNV, and HCV. As our m^6^A change analysis pipeline controls for changes in gene expression, these data should represent true changes in m^6^A modification rather than changes in the expression of m^6^A-modified transcripts. While changes in both m^6^A modification and the expression of m^6^A-modified transcripts are biologically relevant, identifying *bona fide* m^6^A alterations will allow us to understand how m^6^A modification of cellular mRNA is regulated.

Our work reveals that the changes in m^6^A methylation of *RIOK3* and *CRIBP* can be driven by innate immune induction and the cellular response to ER stress, respectively. This suggests that these signals, and likely other infection-induced pathways, can be integrated into differential m^6^A methylation activity and ultimately affect m^6^A modification of cellular mRNAs. Indeed, it is likely that other cellular signaling pathways stimulated by infection can also influence m^6^A modification of cellular transcripts. While changes in the expression of the m^6^A machinery have been shown to affect m^6^A modification during cancer and infection (Barbieri et al., 2017; Li et al., 2017b; Lin et al., 2016a; Rubio et al., 2018; Vu et al., 2017; Winkler et al., 2019), the expression of this machinery did not change with *Flaviviridae* infection, pointing to a different mechanism for altered m^6^A modification. Going forward, identifying the molecular mechanisms through which these signaling pathways lead to differential m^6^A modification during infection will be an important advance in understanding how the cellular m^6^A machinery selects specific sites for modification.

Thus far, our data suggest that the virus-induced m^6^A changes we observed occur in nascent mRNA, which is consistent with the hypothesis that m^6^A is added co-transcriptionally and does not dynamically change at single sites after export to the cytoplasm (Ke et al., 2017). At least three processes could modulate the selective m^6^A modification of specific transcripts during transcription and explain the changes we observed. First, novel interactions of the m^6^A writers METTL3 and METTL14 with viral-induced or stress-regulated RNA-binding proteins could target these writers to specific transcripts and lead to m^6^A epitranscriptomic changes during infection. For example, RBM15/15B and VIRMA can target the m^6^A methyltransferase complex to *Xist* long non-coding RNA or to the 3’ UTRs of mRNA respectively (Patil et al., 2016; Yue et al., 2018). Second, the writers could have differential recruitment to nascent mRNAs by the histone modification H3K36me3 which marks transcriptionally active loci and is known to recruit METTL14 (Huang et al., 2019). Intriguingly, the *CIRBP* locus is marked by H3K36me3 in untreated HepG2 hepatocellular carcinoma cells, and its transcript contains an m^6^A peak at the same site as we identified in Huh7 cells (Huang et al., 2019). This suggests that infection- or ER stress-induced depletion of H3K36me3 marks at the *CIRBP* locus could result in reduced m^6^A modification of *CIRBP* mRNA by METTL3/METTL14. Third, changes in transcription rates, which have been inversely correlated with m^6^A deposition in mRNA, could also contribute to m^6^A modification of specific transcripts during infection (Slobodin et al., 2017). Additionally, viral infection can affect RNA structure in cellular transcripts; it is possible that altered mRNA structure could result in altered m^6^A modification of cellular transcripts during infection (Mizrahi et al., 2018). Perturbing cellular homeostasis by *Flaviviridae* infection therefore has the potential to reveal new insights into how m^6^A modification of cellular transcripts is regulated.

We hypothesize that during viral infection m^6^A regulation of RNA metabolism can lead to rapid, tunable changes in mRNA and protein abundance of host factors. Our work suggests that m^6^A modification promotes translation of *RIOK3* and alternative splicing of *CIRBP*. While m^6^A can affect mRNA nuclear export and stability, *Flaviviridae* infection did not affect these processes for either *RIOK3* or *CIRBP*. m^6^A has also been shown to promote translation of modified mRNAs in multiple contexts by mediating interactions with m^6^A-binding proteins including YTHDF1 (Edupuganti et al., 2017; Han et al., 2019; Huang et al., 2018; Li et al., 2017a; Lin et al., 2016b; Meyer et al., 2015; Shi et al., 2017; Shi et al., 2018; Wang et al., 2019; Wang et al., 2015). We found that increased m^6^A in *RIOK3* mRNA during *Flaviviridae* infection promotes its translation. Interestingly, YTHDF1 also showed increased binding to *RIOK3* during infection. Given its role in recruiting translation factors to modified transcripts and promoting translation under some conditions (Han et al., 2019; Shi et al., 2018; Wang et al., 2019; Wang et al., 2015), YTHDF1 binding to m^6^A in *RIOK3* may allow this transcript to be preferentially translated despite eIF2α phosphorylation and suppression of global translation during infection (Arnaud et al., 2010; Garaigorta and Chisari, 2009; Roth et al., 2017). For *CIRBP*, we found that loss of m^6^A following viral infection led to reduced expression of its long isoform. m^6^A has been shown to regulate splicing by modulating mRNA interactions with several m^6^A-binding splicing factors (Alarcon et al., 2015; Liu et al., 2015; Liu et al., 2017b; Louloupi et al., 2018; Xiao et al., 2016; Ye et al., 2017; Zhao et al., 2014). We hypothesize that the loss of m^6^A in this transcript regulates alternative splicing through changes in the interactions between splicing factors and *CIRBP*. Further investigation into any differences in the roles of the proteins encoded by these two isoforms will help reveal the downstream functional consequences of changes in m^6^A in this gene. How m^6^A regulates the fate of other mRNAs with altered modification also remains unclear, but it is possible that m^6^A post-transcriptionally affects the abundance of their protein products or splicing isoforms, similar to how it regulates *RIOK3* and *CIRBP*.

Importantly, we found that transcripts with altered m^6^A modification during *Flaviviridae* infection encode protein products that can influence the outcome of infection. *RIOK3* expression correlated with the abundance of DENV and ZIKV RNA in a single-cell analysis of infected Huh7 cells, suggesting that it might play a role in infection by these viruses (Zanini et al., 2018). We found that *RIOK3* expression and m^6^A modification was increased with infection by DENV, ZIKV, WNV, and HCV. Further, we found that RIOK3 promoted DENV and ZIKV infection, but inhibited HCV. Interestingly, *RIOK3* has been found to both positively and negatively regulate innate immune responses, by either stimulating the interaction between TBK1 and IRF3 or by phosphorylating and inactivating MDA5 (Feng et al., 2014; Shan et al., 2009; Takashima et al., 2015; Willemsen et al., 2017). The differences in the effects of RIOK3 on DENV, ZIKV, and HCV infection could reflect the different strategies used by these viruses to inhibit host immune responses (Chen et al., 2017; Gack and Diamond, 2016; Gokhale et al., 2014). Further, Willemsen et al. found that while RIOK3 enhanced innate immune activation, it also promoted influenza A virus infection, implying that RIOK3 could have roles in infection beyond innate immunity (Willemsen et al., 2017). Unlike RIOK3, depletion of CIRBP (using siRNA that targets both the small and large isoform) decreased viral replication for DENV, ZIKV, and HCV, suggesting a consistently proviral role. Therefore, during infection, reduction in the long isoform of CIRBP through loss of m^6^A could inhibit infection, suggesting that this loss of m^6^A during infection is part of the host response to infection. CIRBP can modulate the translation of pro-inflammatory factors and have anti-apoptotic effects in response to various stresses (Liao et al., 2017), although the roles of different CIRBP isoforms remains unknown. We also tested whether other transcripts with altered m^6^A following *Flaviviridae* could regulate viral infection. For each virus, approximately half of the factors that we tested that were efficiently depleted showed either proviral or antiviral effects, while in total 86% had roles on any virus. These data suggest that m^6^A itself does not represent a proviral or antiviral mechanism during infection, but rather modulates transcripts that ultimately affect the outcome of infection by different members of the *Flaviviridae* family.

The scale of m^6^A epitranscriptomic changes with virus infection varies greatly among previous reports (Hesser et al., 2018; Lichinchi et al., 2016a; Rubio et al., 2018; Tan et al., 2018; Winkler et al., 2019). Although we identified altered m^6^A in 58 peaks in 51 transcripts during infection, inherent variance in transcript coverage in MeRIP-seq data means that many replicates are necessary for statistically significant detection of m^6^A changes (McIntyre et al., 2019). In particular, this means that our analysis, which used data from three replicates per virus, may underestimate the total number of virus-specific, altered m^6^A peaks. Additionally, we used a more conservative statistical approach than many previous studies to reveal only the most robust peak changes (McIntyre et al., 2019). The changes detected in MeRIP-seq peaks were validated using MeRIP-RT-qPCR; however, these data do not provide the precise ratio of modified to unmodified copies of a transcript or the exact nucleotides that are modified. Biochemical assays like SCARLET or new sequencing methods will be necessary to resolve this question in the future (Liu et al., 2019; Saletore et al., 2012).

In summary, we found that *Flaviviridae* infection leads to m^6^A changes in transcripts that can influence viral infection. We identified innate immune activation and the ER stress response as signals that can modulate m^6^A levels in specific cellular mRNAs. Our work indicates that post-transcriptional regulation of specific transcripts by m^6^A and other RNA modifications can be an important determinant of the outcome of infection. Indeed, viral infection alters the abundance of several other epitranscriptomic modifications on cellular RNA (McIntyre et al., 2018), revealing that we are only at the beginning of our understanding of how m^6^A and other RNA modifications can influence viral infection.

## Acknowledgements

We thank Moonhee Park, Xinhe Yin, Dr. Olga Ilkayeva, Dr. Christopher Nicchitta, and Dr. Heather Vincent for experimental help and advice, colleagues indicated in the methods section for providing reagents, New England Biolabs for donating anti-m^6^A antibodies, the Metabolomics Core at the Duke Molecular Physiology Institute, the Duke Functional Genomics Core Facility, the Epigenomics Core and the Scientific Computing Unit at Weill Cornell, and Dr. Kate Meyer, Danielle Dauphars, and members of the Horner lab for discussion and reading of this manuscript. This work was supported by funds from the Burroughs Wellcome Fund (S.M.H.) and the National Institutes of Health: R01AI125416, R21AI129851 (S.M.H. and C.E.M.), 5P30AI064518 (S.M.H.), R01MH117406 (C.E.M.), R03HL135475 (C.L.H), and R21AI129431 (H.M.L). Other funding sources include: the American Heart Association (N.S.G. Pre-doctoral Fellowship, 17PRE33670017), the National Science and Engineering Research Council of Canada (A.B.R.M. PGS-D funding), The Bert L. and N. Kuggie Vallee Foundation, the WorldQuant Foundation, the Pershing Square Sohn Cancer Research Alliance, NASA (NNX14AH50G), and startup finds from the University of North Carolina Lineberger Comprehensive Cancer Center (H.M.L. and M.D.M).

## Author contributions

Conceptualization: N.S.G., A.B.R.M., C.E.M., and S.M.H. Investigation: N.S.G., A.B.R.M., C.L.H., H.M.L., and M.D.M. Formal analysis: A.B.R.M. and N.S.G. Software: A.B.R.M. Visualization: N.S.G. and A.B.R.M. Writing – original draft: N.S.G., A.B.R.M., and S.M.H. Writing – review and editing: N.S.G., A.B.R.M., C.L.H., H.M.L., M.D.M., C.E.M., and S.M.H. Funding acquisition: N.S.G., A.B.R.M., C.L.H., H.M.L., C.E.M., and S.M.H.

## Methods

### Cell culture, viral stocks, and viral infection

Huh7 and Huh-7.5 cells (gift of Dr. Michael Gale Jr., University of Washington (Sumpter et al., 2005)), 293T cells (ATCC: CRL-3216) Vero cells (ATCC: CCL-81), C6/36 (ATCC: CRL-1660) were grown in Dulbecco’s modification of Eagle’s medium (DMEM; Mediatech) supplemented with 10% fetal bovine serum (HyClone), 25 mM HEPES (Thermo Fisher), and 1X non-essential amino acids (Thermo Fisher), referred to as complete DMEM (cDMEM). Huh7 and Huh-7.5 cells were verified using the Promega GenePrint STR kit (DNA Analysis Facility, Duke University), and cells were verified as mycoplasma free by the LookOut Mycoplasma PCR detection kit (Sigma-Aldrich). Infectious stocks of a cell culture-adapted strain of genotype 2A JFH1 HCV were generated and titered in Huh-7.5 cells by focus-forming assay (FFA), as described (Aligeti et al., 2015). DENV2-NGC (Sessions et al., 2009), ZIKV-PR2015 (Quicke et al., 2016), and WNV-NY2000 (Diamond et al., 2003) stocks were prepared in C6/36 insect cells and titered in Vero cells, as described. For viral infections, cells were incubated in a low volume of cDMEM containing virus at a multiplicity of infection (MOI) of one for 2-3 hours (except when otherwise stated), following which cDMEM was replenished. Cells were infected for 48 hours unless otherwise described. To quantify virus, cellular supernatants were analyzed by FFA.

### MeRIP-seq and MeRIP-RT-qPCR

#### Sample preparation

Huh7 cells seeded in 15 cm plates were infected with DENV, ZIKV, WNV, or HCV (MOI 1) or left uninfected (mock-infected). At 48 hours post-infection, total RNA was extracted using TRIzol (Thermo Fisher) and treated with TURBO DNase I (Thermo Fisher). mRNA was purified from 200 µg total RNA from each sample using the Dynabeads mRNA purification kit (Thermo Fisher) and concentrated by ethanol precipitation. mRNA was fragmented using the RNA Fragmentation Reagent (Thermo Fisher) for 15 minutes and purified by ethanol precipitation. MeRIP was performed using EpiMark *N6*-methyladenosine Enrichment kit (NEB) according to the manufacturer’s recommendations with the following modifications. Briefly, 25 µL Protein G Dynabeads (Thermo Fisher) per sample were washed three times in MeRIP buffer (150 mM NaCl, 10 mM Tris-HCl [pH 7.5], 0.1% NP-40), and incubated with 1 µL anti-m^6^A antibody for 2 hours at 4°C with rotation. After washing three times with MeRIP buffer, anti-m^6^A conjugated beads were incubated with purified mRNA with rotation at 4°C overnight in 300 µL MeRIP buffer with 1 µL RNase inhibitor (recombinant RNasin; Promega). 10% of the mRNA sample was saved as the input fraction. Beads were then washed twice with 500 µL MeRIP buffer, twice with low salt wash buffer (50 mM NaCl, 10 mM Tris-HCl [pH 7.5], 0.1% NP-40), twice with high salt wash buffer (500 mM NaCl, 10 mM Tris-HCl [pH 7.5], 0.1% NP-40), and once again with MeRIP buffer. m^6^A-modified RNA was eluted twice in 100 µL of MeRIP buffer containing 5 mM m^6^A salt (Santa Cruz Biotechnology) for 30 minutes at 4°C with rotation. Eluates were pooled and concentrated by ethanol purification. RNA-seq libraries were prepared from both eluate and 10% input mRNA using the TruSeq mRNA library prep kit (Illumina), subjected to quality control (MultiQC), and sequenced on the HiSeq 4000 instrument.

For MeRIP-RT-qPCR, total RNA was harvested from cells with the indicated treatments in 10 cm plates or 6-well plates. For ER-stress induction, cells seeded in 6-well plates were treated with 500 nM thapsigargin (Tocris) for 16 hours. HCV PAMP was prepared by *in vitro* transcription, as described (Beachboard et al., 2019; Saito et al., 2008). 2.5 µg of HCV PAMP RNA was transfected into cells seeded in 6-well plates using the Mirus mRNA transfection kit. 8 hours later, RNA was extracted and MeRIP-RT-qPCR was performed like MeRIP-seq with some differences. Specifically, total RNA was prepared from cells using TRIzol, and diluted to equivalent concentrations. Then, 20-50 µg total RNA was fragmented for 3 minutes, purified by ethanol precipitation, and resuspended in 30 µL water. 0.1 fmol of positive control (m^6^A-modified *Gaussia* luciferase RNA) and negative control (unmodified *Cypridina* luciferase RNA) spike-ins supplied with the EpiMark *N6*-methyladenosine Enrichment kit were added to each sample. Following MeRIP as described above, eluates were concentrated by ethanol precipitation. 1 µL input and the entire IP fractions were reverse transcribed using the iScript cDNA synthesis kit (BioRad) and subjected to RT-qPCR. Primer sequences are supplied in Table S3. Relative m^6^A level for each transcript was calculated as the percent of input in each condition normalized to that of the respective positive control spike-in. Fold change of enrichment was calculated with mock samples normalized to 1.

#### Data analysis

Reads were aligned using STAR (Dobin et al., 2013) to the human reference genome (hg38), combined with the appropriate virus genome for each infected sample. Differential gene expression between infected and uninfected samples was compared using DESeq2 (Love et al., 2014). UpSet plots of the intersects between genes regulated with individual viruses were generated using UpSetR (Conway et al., 2017). Gene ontology for RNA-seq changes in Figure S1D was analyzed using gProfiler, with redundant GO terms collapsed using REVIGO (Reimand et al., 2016; Supek et al., 2011). For gProfiler, upregulated genes with Log_2_FC ≥ 2 and adjusted p-value < 0.05 with all viruses were considered. There were very few consistently downregulated genes at Log_2_FC ≤ −2 (particularly for ZIKV), so we expanded our set to genes with smaller Log_2_FC ≤ −0.5, downregulated by DENV, HCV, and WNV infection. For REVIGO, we allowed similarity of up to 0.5, with semantic similarity calculated using SimRel. Adjusted p-values were provided for the REVIGO calculations. Gene set enrichment analyses using fgsea in R showed similar differentially regulated pathways as gProfiler (Sergushichev, 2016). “Infection-annotated” genes and peaks were summarized for Figure 1B based on gene inclusion in “Infectious disease”, “Unfolded Protein Response (UPR)”, “Interferon Signaling”, and “Innate Immune System” Reactome pathways from fgsea.

We called m^6^A peaks from MeRIP-seq using MACS2 (Zhang et al., 2008) and used all peaks detected in at least two replicates for further analysis. Motif enrichment was calculated using HOMER for Figure 1C (Heinz et al., 2010). Metagene plots for methylated DRACH motifs were plotted using a custom script. DRACH motifs were considered methylated if detected under m^6^A peaks in at least 2 biological replicates. Relative positions of m^6^A peaks within genes are based on the transcripts with the highest mean coverage per gene, as calculated with kallisto (Bray et al., 2016).

We identified m^6^A peaks changes using a generalized linear model (adapted from (Park et al., 2014)), and the QNB program (Liu et al., 2017a). In brief (see Park et al., 2014 or McIntyre et al., 2019 for more details), a generalized linear model following the equation

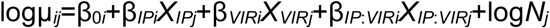

was fit with the following parameters for each peak i and sample j: X_IP_ = 1 for immunoprecipitated samples and 0 for input samples, and X_VIR_ = 1 for infected samples and 0 for mock. A library size parameter was included for normalization (N) with edgeR (Robinson et al., 2010). The full model was compared to a reduced model without the infection:IP interaction term using a likelihood ratio test of the difference between deviances, implemented through DESeq2 (Love et al., 2014) or edgeR. To control for changes in gene expression, changes in gene expression were subtracted from changes in IP peak reads for significantly modified peaks from DESeq2, edgeR, and QNB, with a threshold for absolute difference in Log_2_ fold change of ≥ 1. Significant peaks were further filtered for location within exons, DRACH motif content, and mean input read counts of ≥ 10 to produce the final set of 58 peak changes.

Peaks of interest were plotted for visual evaluation using CovFuzze (https://github.com/al-mcintyre/CovFuzze) (Imam et al., 2018).

### RT-qPCR

The iScript cDNA synthesis kit (Bio-Rad) was used for reverse transcription of total RNA samples. RT-qPCR was performed using the Applied Biosystems QuantStudio 6 Flex real-time PCR instrument. To measure relative abundance of *CIRBP* isoforms, total RNA was reverse transcribed with the Superscript III enzyme (Invitrogen) using a gene specific primer. RT-qPCR was performed using specific primers that detect *CIRBP* isoforms. The expression of each isoform was normalized to invariant region of *CIRBP*. Primer sequences are provided in Table S3.

### Immunoblotting

Cell lysates were prepared in a modified RIPA buffer (10 mM Tris [pH 7.5], 150 mM NaCl, 0.5% sodium deoxycholate, and 1% Triton X-100) supplemented with protease inhibitor cocktail (Sigma-Aldrich) and phosphatase inhibitor cocktail II (Millipore), and clarified by centrifugation. Protein concentration was determined by Bradford assay (Bio-Rad). 5-15 µg of protein was resolved by SDS/PAGE and transferred to nitrocellulose membranes using the Trans-Blot Turbo System (Bio-Rad). Membranes were blocked in 5% milk in phosphate buffered saline with 0.1% Tween (PBS-T) and incubated with the relevant primary antibodies. After washing three times with PBS-T, membranes were incubated with species-specific horseradish peroxidase-conjugated antibodies (Jackson ImmunoResearch, 1:5000) or fluorescent antibodies (LI-COR, IRDye 800, 1:5000). Chemiluminescence (Clarity ECL, Bio-Rad) or fluorescence was detected on a LI-COR Odyssey Fc instrument and analyzed using the ImageStudio software. The following antibodies were used for immunoblot: anti-METTL3 (Novus Biologicals, 1:1000), anti-METTL14 (Sigma-Aldrich, 1:5000), anti-FTO (Abcam, 1:1000), anti-YTHDF1 (Proteintech, 1:1000), anti-YTHDF2 (Proteintech, 1:1000), anti-YTHDF3 (Sigma-Aldrich, 1:1000), anti-ALKBH5 (Sigma-Aldrich, 1:1000), anti-WTAP (Proteintech, 1:1000) anti-FLAG M2 (Sigma-Aldrich, 1:5000), anti-tubulin (Sigma-Aldrich, 1:5000), anti-HCV NS5A (clone 9E10, gift of Charles Rice, Rockefeller University (Lindenbach et al., 2005), 1:1000), anti-RIOK3 (Proteintech, 1:1000), anti-CIRBP (Proteintech 1:1000), anti-DENV NS3 (GeneTex, 1:1000), anti-ZIKV NS3 (GeneTex, 1:1000), anti-HCV NS4A (Genscript custom (Horner et al., 2011)), 1:1000), anti-eIF2α (Cell Signaling, 1: 1000), anti-phospho-eIF2α (Cell Signaling, 1:1000), anti-GADD34 (Proteintech, 1:1000), anti-HSPA5 (Cell Signaling, 1:1000), anti-H2A.X (Cell Signaling, 1:1000), anti-U170K serum (gift of Dr. Jack Keene, Duke University, (Query and Keene, 1987), 1:1000)

### FLAG-YTHDF RNA immunoprecipitation

Generation of Huh7 cells stably expressing FLAG-GFP or FLAG-YTHDF1 was described previously (Gokhale et al., 2016). Cells seeded in 6-well plates were infected with DENV, ZIKV, or HCV (MOI 1). At 48 hours post-infection cells were harvested by trypsinization and lysed in polysome lysis buffer (100 mM KCl, 5 mM MgCl_2_, 10 mM HEPES [pH 7.0], 0.5% NP-40), supplemented with protease inhibitor cocktail (Sigma-Aldrich) and RNase inhibitor (RNasin), and cleared by centrifugation. Protein was quantified by Bradford assay, and 200 µg ribonucleoprotein complexes were immunoprecipitated with M2 anti-FLAG conjugated magnetic beads (Sigma-Aldrich) overnight at 4°C with rotation in NT2 buffer (50 mM Tris-HCl [pH 7.5], 150 mM NaCl, 1 mM MgCl_2_, 0.05% NP-40). Beads were washed five times in ice-cold NT2 buffer. Protein for immunoblotting was eluted from ten percent of beads by boiling in 2X Laemmli sample buffer (Bio-Rad). RNA was extracted from ninety percent of beads using TRIzol reagent (Thermo Fisher). Equal volumes of eluted RNA were used for cDNA synthesis, quantified by RT-qPCR, and normalized to RNA levels in input samples. Fold enrichment was calculated with FLAG-GFP and mock samples set as 1.

### siRNA treatment and viral infectivity assays

Cells seeded in 24-well plates were transfected with siRNA against intended targets (Qiagen, sequences provided in Table S3) using Lipofectamine RNAiMAX (Thermo Fisher) according to the manufacturer’s recommendation. At 24 hours post-transfection, cells were infected with DENV, ZIKV, and HCV (MOI 0.2). At 48 (targeted siRNA screen) or 72 (RIOK3 and CIRBP depletion) hours post-infection, virus titer in the supernatant was measured by FFA. Serial dilutions of supernatants were used to infect naïve Vero (DENV and ZIKV) or Huh-7.5 (HCV) cells in triplicate wells of a 48-well plate. At 72 hours post-infection, cells were fixed in cold 1:1 methanol:acetone and immunostained with 4G2 antibody purified in the lab from a hybridoma (for DENV and ZIKV, 1:2000), or anti-HCV NS5A (1:2000). Following binding of horseradish peroxidase conjugated secondary antibody (1:1000; Jackson ImmunoResearch), infected foci were visualized with the VIP Peroxidase Substrate Kit (Vector Laboratories) and counted at 40X magnification. Titer was calculated using the following formula: (dilution factor × number of foci × 1000) / volume of infection (μl), resulting in units of focus forming units / mL (FFU/mL). Depletion of siRNA targets was confirmed by RT-qPCR (primer sequences in Table S3). Cellular viability after siRNA treatment was measured by the Cell-Titer Glo assay (Promega) according to the manufacturer’s recommendation.

### Quantification of infection by immunofluorescence

To measure percent of cells infected following viral infection, Huh7 cells seeded in 96-well plates were infected with DENV, ZIKV, WNV, or HCV (MOI 1). Cells were fixed in cold 1:1 methanol:acetone at the indicated hours post-infection, and immunostained with 4G2 antibody (DENV, ZIKV, WNV) or anti-HCV NS5A. Following binding of AlexaFluor 488-conjugated secondary antibody (Thermo Fisher) and nuclear staining with Hoechst (Thermo Fisher), cells were imaged using the Cellomics Arrayscan VTI robotic microscope. The percentage of infected cells was determined by measuring cells stained for viral antigen relative to the total number of nuclei.

### Cell fractionation

Fractionation of cells to isolate chromatin-associated RNA was performed as described (Ke et al., 2017). Briefly, cells were collected from 10 cm plates by trypsinization, lysed in 200 µL cytoplasmic lysis buffer (10 mM Tris-HCL [pH 7.4], 150 mM NaCl, 0.15% NP-40) on ice for 5 minutes, and passed through 500 µl 24% sucrose cushion by centrifugation at 12000 xG for 10 minutes at 4°C. The supernatant (cytoplasmic fraction) was then removed and the nuclear pellet was rinsed twice with cold phosphate buffered saline (PBS). The nuclear pellet was resuspended in 100 µL ice cold glycerol buffer (20 mM Tris-HCL [pH 7.4], 75 mM NaCl, 0.5 mM EDTA, 1 mM DTT, 125 µM PMSF, 50% glycerol). 100 µL nuclear lysis buffer (10 mM HEPES [pH 7.4], 1 mM DTT, 7.5 mM MgCl_2_, 0.2 mM EDTA, 300 mM NaCl, 1 M urea, 1% NP-40) was added to the suspension, followed by brief vortexing, and incubation on ice for 2 minutes. Samples were centrifuged for 2 minutes at 4°C at 12000 xG and the supernatant (nuclear fraction) was removed. The chromatin pellet was rinsed twice with cold PBS, resuspended in 50 µL DNase I buffer with 2 U Turbo DNase I (Invitrogen), and incubated at 37°C for 30 minutes. RNA was then extracted from the chromatin fraction using TRIzol reagent and subjected to MeRIP-RT-qPCR. The cytoplasmic, nuclear, and chromatin fractions were subjected to immunoblotting to analyze fractionation.

For nuclear/cytoplasmic fractionation to investigate mRNA export, uninfected and infected (MOI 1) cells grown in 10 cm plates were harvested by trypsinization and lysed in 200 µL lysis buffer (10mM Tris-HCl [pH 7.4], 140 mM NaCl, 1.5 mM MgCl_2_, 10 mM EDTA, 0.5% NP-40) on ice for 5 minutes. Following centrifugation at 12000 xG at 4°C for 5 minutes, the supernatant (cytoplasmic fraction) was collected, and the nuclear pellet was rinsed twice with lysis buffer. RNA was extracted from cytoplasmic and nuclear pellets using TRIzol reagent and analyzed by RT-qPCR.

### Measurement of RNA stability

Cells plated in 24-well plates were infected with the indicated virus (MOI 1). At 36 hours post-infection, media was changed to cDMEM containing 1 µM Actinomycin D (Sigma-Aldrich). RNA was extracted from cells at the indicated time points post-treatment using TRIzol reagent and analyzed by RT-qPCR. Data were normalized as the percent of RNA remaining at each time point after treatment, relative to that at the time of treatment.

### Cloning of *RIOK3* and *CIRBP* and generation of stable cell lines

All primer sequences used for cloning are provided in Table S3. *RIOK3* (NM_003831.4) and both long (NM_001300829) and short (NM_001280) isoforms of *CIRBP* were cloned by PCR (HiFi PCR premix, Clontech) from cDNA from Huh7 cells prepared with the Superscript III RT kit (Thermo Fisher) using the oligo(dT)_20_ primer. PCR products were inserted into pLEX-FLAG lentiviral vector between the *NotI* and *XhoI* sites using the InFusion HD cloning kit (Takara Bio) to generate constructs with N-terminal FLAG tags. Lentivirus was produced from 293T cells transfected with pLEX vectors and packaging plasmids psPAX2 and pMD2.G (provided by Duke Functional Genomics Facility).

Huh7 cells were transduced by these lentiviruses and stable cell lines expressing FLAG-RIOK3, FLAG-CIRBP-S, and FLAG-CIRBP-L were selected using puromycin (2 µg/mL). Single cell clones were obtained by serial dilution and verified by immunoblotting. Cell lines were maintained in cDMEM containing 1 µg/mL puromycin.

### Reporter cloning and luciferase assays

All primer and gBlock sequences are provided in Table S3. To generate m^6^A-null *RIOK3* reporters, the *Renilla* and Firefly luciferase genes in psiCheck2 plasmid (Promega) were first replaced by constructs with synonymous mutations in putative m^6^A sites (obtained as IDT gBlocks). The wild type *RIOK3* 3’ UTR was cloned from Huh7 cDNA (NM_003831.4) and inserted after the m^6^A-null *Renilla* luciferase gene in the multiple cloning site of psiCheck2 between *XhoI* and *NotI* using the InFusion HD kit. m^6^A-mut *RIOK3* 3’ UTR (in which all putative m^6^A sites were mutated from A to T) was obtained as a gBlock and also inserted between these restriction sites. WT and m^6^A-mut *RIOK3* reporter plasmids along with the pcDNA-Blast plasmid (Kennedy et al., 2015) were linearized using *BamHI* and *BglII* respectively, purified by ethanol precipitation and co-transfected into Huh7 cells in 6-well plates (90 ng reporter, 10 ng pcDNA-Blast) using FuGENE 6 transfection reagent (Promega). Cells were selected with blasticidin (0.2 μg/mL) and single cell clones stably expressing WT and m^6^A-mut reporters were isolated. For MeRIP-RT-qPCR of reporter RNA, WT and m^6^A-mut expressing cells were plated in 6-well plates, infected with the indicated virus (MOI 1), and RNA was extracted using TRIzol at 48 hours post-infection. Following MeRIP as described, RT-qPCR was performed to discriminate reporter RNA using a forward primer within the *Renilla* luciferase gene and a reverse primer in the *RIOK3* 3’ UTR. For luciferase assays, WT and m^6^A-mut expressing cells in 24-well plates were infected with the indicated virus (MOI 1) and dual luciferase assay (Promega) was performed at 48 hours post-infection according to the manufacturer’s instructions. Data was normalized as the value of Renilla luminescence divided by Firefly luminescence, and values for mock-infected cells were set as 1.

To generate *CIRBP* splicing reporters, *CIRBP* exon 5 – 3’ UTR (Hg38;chr19:127553-1273172) was amplified by PCR from genomic DNA. A fragment of m^6^A-null *Renilla* luciferase beyond the *NruI* site and up to the stop codon was amplified by PCR with overlapping ends with *Renilla* luciferase (5’; before the *NruI* site) and the *CIRBP* fragment (3’). These fragments were inserted into *NruI*-*XhoI* digested psiCheck2 m^6^A-null plasmid using the InFusion HD kit. m^6^A-mut *CIRBP* reporter was generated by mutating the essential C in the m^6^A site synonymously to T using two rounds of site-directed mutagenesis with the QuikChange Lightning kit (Agilent).

### ^35^S labeled immunoprecipitation for nascent translation

Huh7 cells seeded in 10 cm plates were infected with DENV (MOI 1) or left uninfected. At 45 hours post-infection, media was removed and 3 mL warm methionine/cysteine-free DMEM was added to plates. After 15 minutes of incubation, 3 mL methione/cysteine-free DMEM containing 100 mCi ^35^S (Perkin Elmer) was added. Cells were harvested at 3 hours post-treatment and lysed in RIPA buffer. 300 µg protein was incubated with 4 µg anti-RIOK3 antibody (Proteintech) or normal rabbit IgG (Cell Signaling) in 300 µL RIPA buffer overnight at 4°C with rotation. Antibody-protein complexes were then incubated with 40 µL pre-washed protein G Dynabeads (Thermo Fisher) for 2 hours. Protein was eluted from beads in 2X Laemmli buffer. Eluates were resolved by SDS/PAGE. Gels were fixed in solution containing 50% methanol and 10% acetic acid, dried, and subjected to autoradiography on film.

### LC-MS/MS for m^6^A/A determination

mRNA was purified from 200 µg total RNA extracted from uninfected and infected Huh7 cells (MOI 1, 48 hours post-infection) using one round of polyA selection (Dynabeads mRNA purification kit; Thermo Fisher) and one round of rRNA depletion (NEBNext rRNA depletion kit, NEB). After ethanol precipitation, purified mRNA was digested into mononucleotides with nuclease P1 (Sigma-Aldrich, 2 U) in buffer containing 25 mM NaCl and 2.5 mM ZnCl_2_ for 2 hours at 37°C, followed by incubation with Antarctic Phosphatase (NEB, 5 U) for an additional 2 hours at 37°C. Nucleosides were separated and quantified using UPLC-MS/MS as previously described, except acetic acid was used in place of formic acid (Basanta-Sanchez et al., 2016).

### Data Availability

All raw data from MeRIP-seq analysis of uninfected and infected Huh7 cells are available through GEO (accession number: GSE130891).

