## Supplemental Information for "Altered m^6^A modification of specific cellular transcripts affects *Flaviviridae* infection"

**Table of contents:**

Figure S1: Related to Figure 1.

Figure S2: Related to Figure 2.

Figure S3: Related to Figure 3.

Figure S4: Related to Figure 4.

Figure S5: Related to Figure 5.

**Figure S1: Related to Figure 1.**

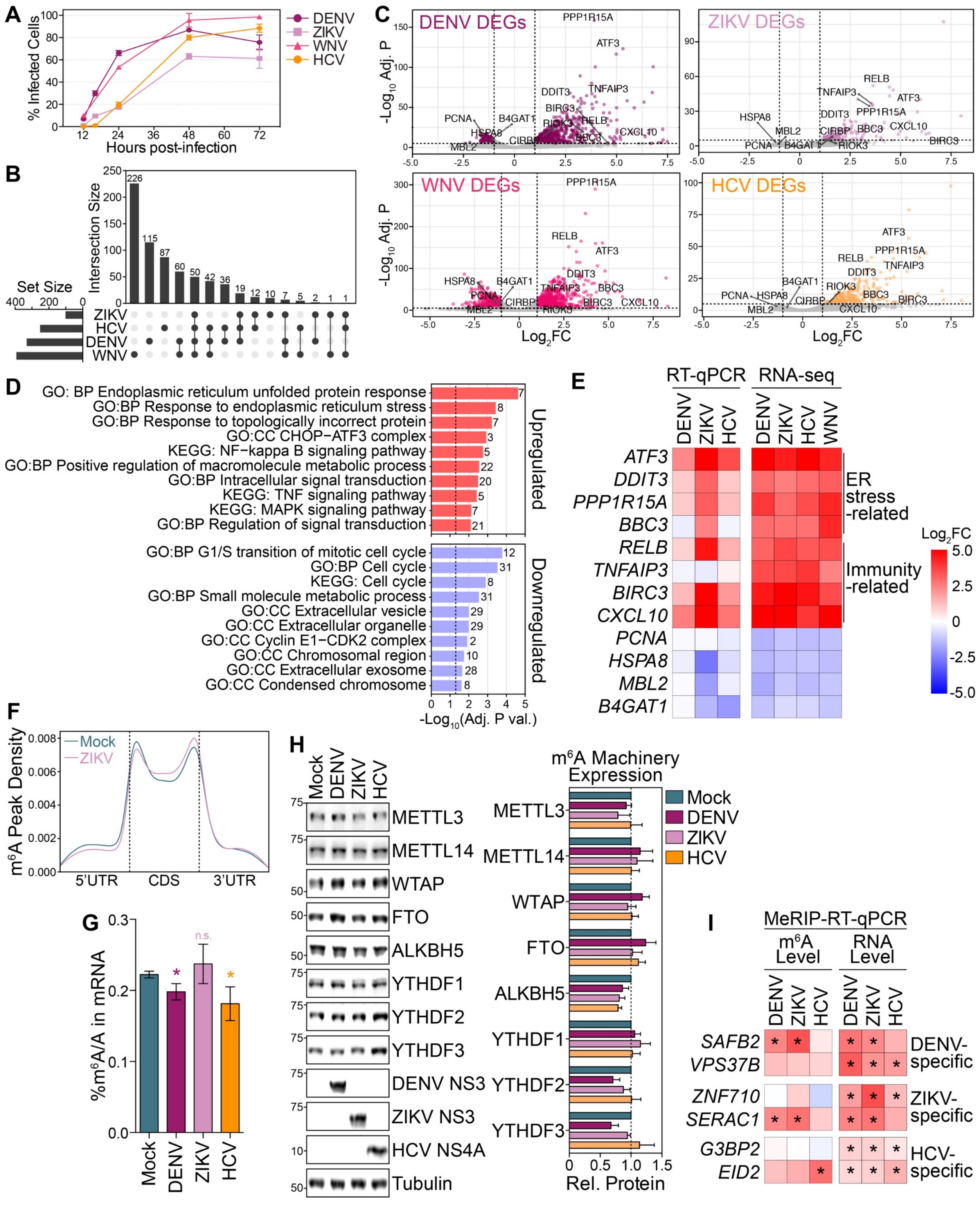

**Figure S1: Related to Figure 1. (A)** Percent of Huh7 cells infected with DENV, ZIKV, WNV, and HCV at the indicated hours post-infection as determined by immunostaining of viral antigen and nuclei. >5000 cells counted for each condition. **(B)** UpSet plot showing the number of differentially expressed genes (DEGs, $|Log_2FC| \geq 2$, adjusted $p < 0.05$) and those in common with DENV, ZIKV, WNV, and HCV infection as determined by RNA-seq analysis of input fractions from MeRIP-seq using DESeq2. **(C)** Volcano plots of DEGs following infection by the indicated virus. Colored dots represent significant DEGs ($|Log_2FC| \geq 2$, adjusted $p < 0.05$). Example genes validated by RT-qPCR in (E) are named. **(D)** Pathway enrichment (using gProfiler) for genes upregulated (top; red color) or downregulated (bottom; blue color) by DENV, ZIKV, WNV, and HCV infection. Few genes were significantly downregulated with ZIKV infection and were excluded from this analysis. Numbers refer to the number of genes in each category. **(E)** Heatmap of $Log_2FC$ in RNA levels for RT-qPCR validation at 48 hpi (left) of DEGs co-regulated by infection in Huh7 cells, as determined by RNA-seq (right). Values in RT-qPCR heatmaps were normalized to *GAPDH* and represent the mean of 3 biological replicates. **(F)** Metagene plot of methylated DRACH motifs across transcripts in mock- and ZIKV-infected cells in 293T cells reanalyzed from (Lichinchi et al., 2016b). DRACH motifs were considered methylated if they fell under a peak detected in at least two replicates. **(G)** LC-MS/MS quantification of $m^6A/A$ ratios in purified mRNA from mock- and virus-infected (48 hpi) Huh7 cells. Values are the mean ± SEM of three experiments. * $p \leq 0.05$, by unpaired Student's t test. n.s. = not significant. **(H)** (Left) Representative immunoblot of protein expression of the cellular $m^6A$ machinery in mock- and virus-infected (48 hpi) Huh7 cells. (Right) Quantification of immunoblot analysis of the cellular $m^6A$ machinery relative to tubulin. Values are the mean ± SEM of 4 biological replicates. **(I)** (Left) MeRIP-RT-qPCR analysis of relative $m^6A$ level of transcripts identified as having altered $m^6A$ modification with the indicated virus in DENV, ZIKV, and HCV-infected (48 hpi) Huh7 cells. (Right) RNA expression of these transcripts relative to *GAPDH* (right). Values in heatmaps are the mean of 3 independent experiments. * $p < 0.05$, by unpaired Student's t test.

**Figure S2: Related to Figure 2.**

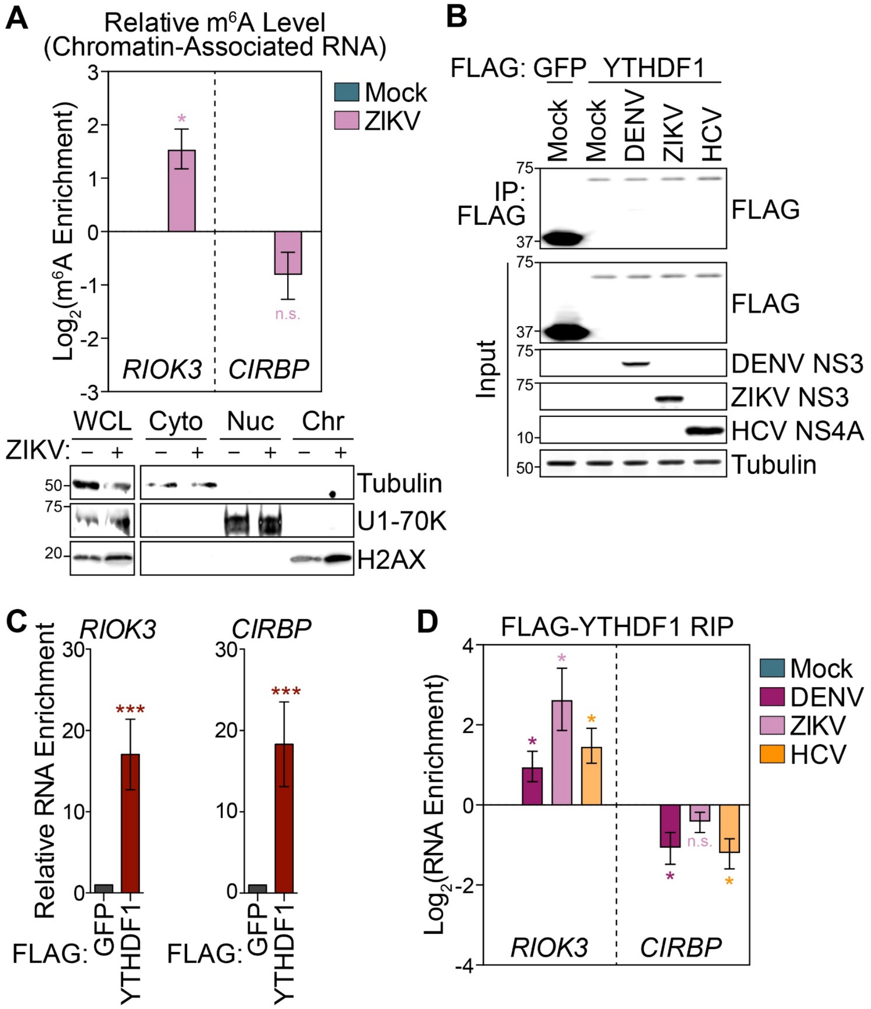


**Figure S2. Related to Figure 2. (A)** (Top) MeRIP-RT-qPCR analysis of relative m$^6$A level of chromatin-associated *RIOK3* and *CIRBP* RNA in mock- and ZIKV-infected (48 hpi) Huh7 cells. (Bottom) Immunoblot analysis of whole cell lysate (WCL), cytoplasmic (Cyto), nuclear (Nuc), and chromatin (Chr) fractions. **(B)** Representative immunoblot of FLAG-immunoprecipitated (IP) and input fractions used for RT-qPCR analysis in (C) and (D). **(C)** RT-qPCR analysis of enrichment of *RIOK3* and *CIRBP* RNA by immunoprecipitation of FLAG-YTHDF1 compared to FLAG-GFP in uninfected Huh7 cells stably expressing these constructs. **(D)** RT-qPCR analysis of enrichment of *RIOK3* and *CIRBP* RNA by immunoprecipitation of FLAG-YTHDF1 in mock- or virus-infected (48 hpi) Huh7 cells expressing FLAG-YTHDF1. Relative enrichment was calculated as the percent of input for each sample relative to that in mock-infected cells. For (A, C, and D), values are the mean ± SEM of 2-3 biological replicates. * $p < 0.05$, *** $p < 0.001$ by unpaired Student's t test. n.s. = not significant.

**Figure S3: Related to Figure 3.**

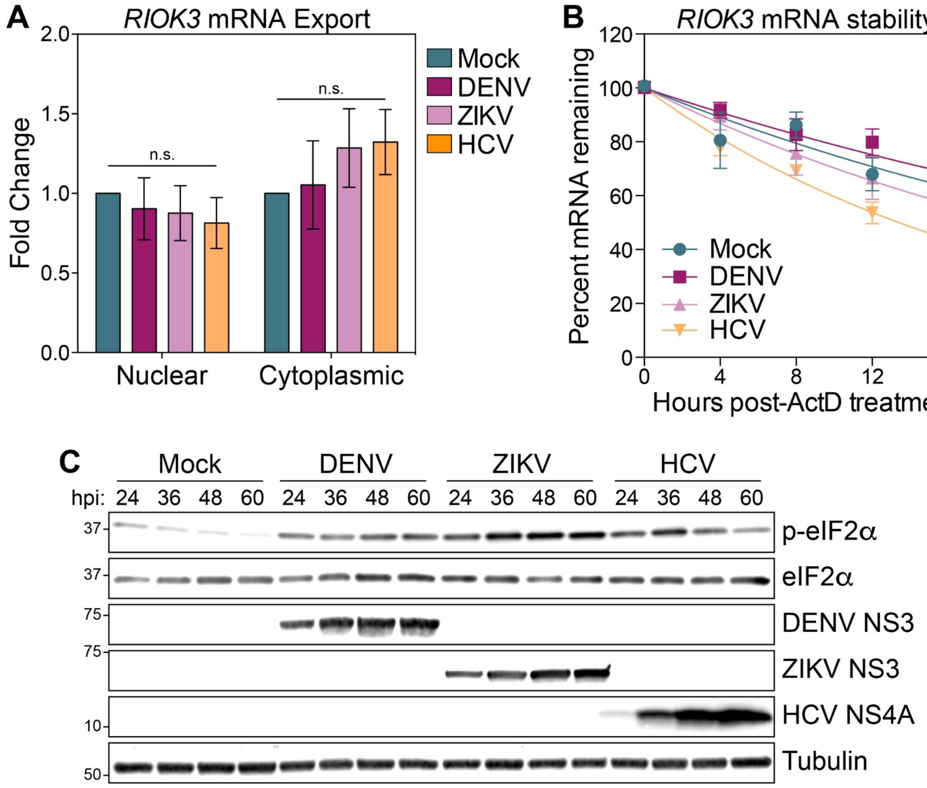


**Figure S3: Related to Figure 3. (A)** RT-qPCR quantification of relative *RIOK3* RNA in nuclear and cytoplasmic fractions in mock- and virus-infected (48 hpi) Huh7 cells. **(B)** Measurement of *RIOK3* RNA in mock- and virus-infected Huh7 cells. At 36 hpi, cell culture media was replaced with media containing actinomycin D (ActD). RNA was harvested cells at the indicated times post-treatment and subjected to RT-qPCR to determine remaining relative RNA levels. **(C)** Immunoblot analysis of p-eIF2α and eIF2α levels in mock- and virus-infected Huh7 cells at the indicated time points. Data are representative of 3 biological replicates. For (A and B), values represent mean ± SEM from 3 biological replicates. n.s. = not significant.

**Figure S4: Related to Figure 4.**

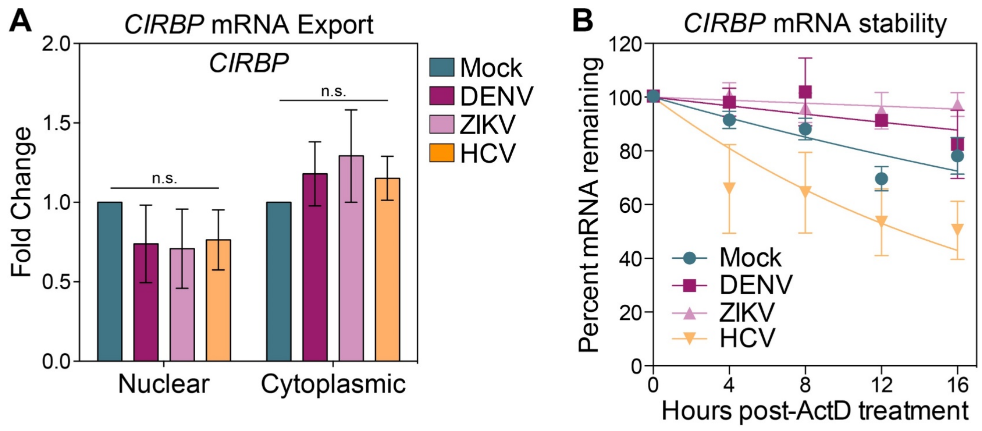


**Figure S4: Related to Figure 4. (A)** RT-qPCR quantification of relative *CIRBP* RNA in nuclear and cytoplasmic fractions in mock- and virus-infected (48 hpi) Huh7 cells. **(B)** Measurement of *CIRBP* RNA in mock- and virus-infected Huh7 cells. At 36 hpi, cell culture media was replaced with media containing ActD. RNA was harvested cells at the indicated times post-treatment and subjected to RT-qPCR to determine remaining relative RNA levels.

**Figure S5: Related to Figure 5.**

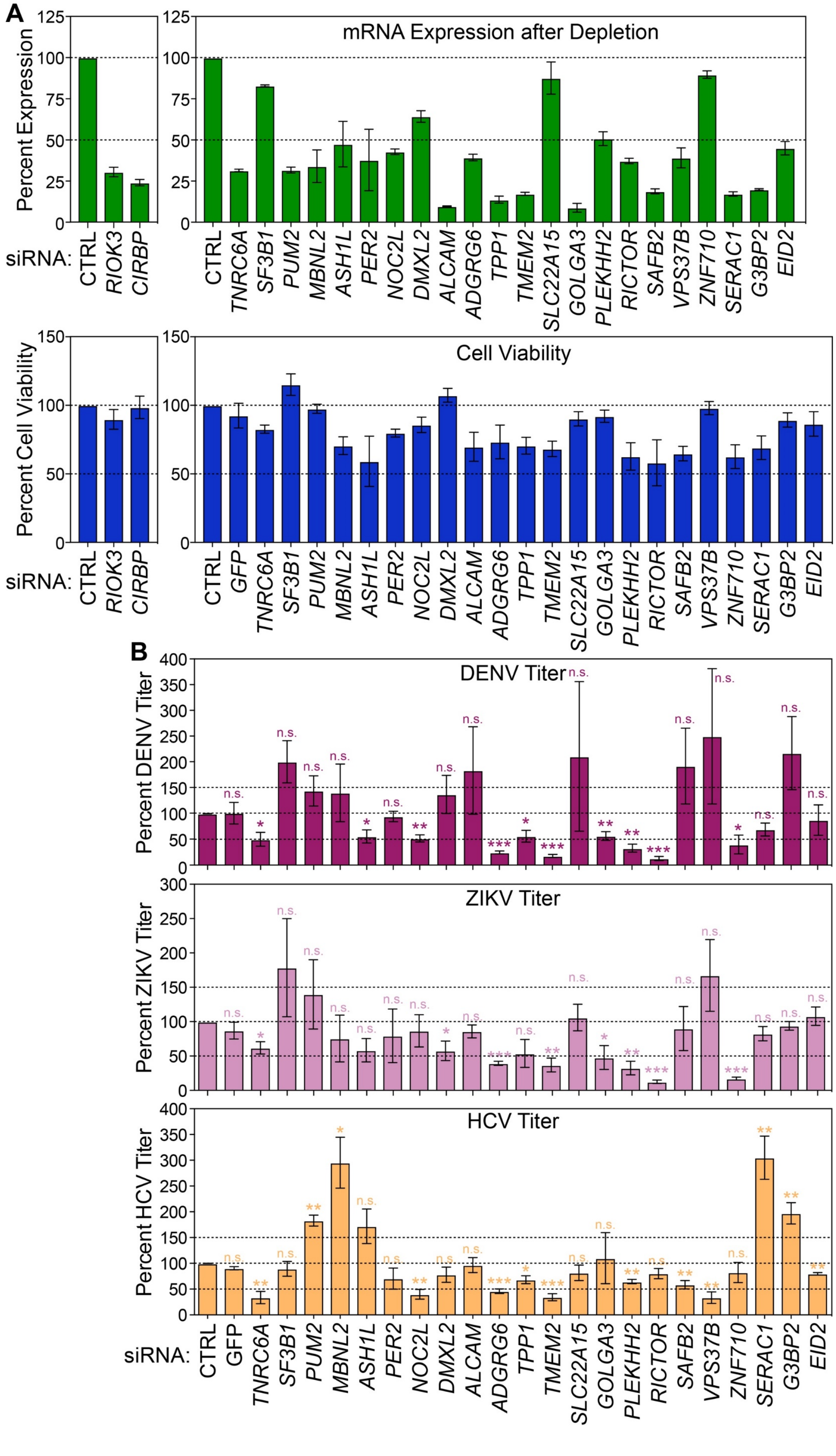

**Figure S5: Related to Figure 5. (A)** (Top) RT-qPCR of the indicated transcripts at 72 hours post-transfection of siRNA compared to non-targeting control (CTRL) siRNA. All values are relative to *GAPDH* expression. (Bottom) Cell viability measured after siRNA depletion of indicated transcripts at 72 hours post-transfection of siRNA relative to that of siCTRL. Expression or cell viability in cells treated with siCTRL was set at 100%. **(B)** FFA of supernatants harvested at 48 hpi from Huh7 cells treated with the indicated siRNAs and infected with DENV (top), ZIKV (middle), or HCV (bottom). Viral titer in siCTRL treated cells was set at 100%. All viral infections for experiments in this figure were performed at a multiplicity of infection of 0.2. All values are the mean ± SEM of three biological replicates. * $p < 0.05$, ** $p < 0.01$, *** $p < 0.001$ by unpaired Student's t test. n.s. = not significant.
